# Mechanobiology-guided drug repurposing identifies budesonide as an inhibitor of stiffness-induced PDAC aggressiveness

**DOI:** 10.64898/2026.07.17.739125

**Authors:** Simone Mas, Marco Cristiano, Eduardo Ibello, Alessandro Avallone, Crescenzo Frascogna, Bruno Sainz, Enza Lonardo, Lucia Altucci, Gilda Cobellis, Eduardo Jorge Patriarca, Paolo Netti, Gabriella Minchiotti, Valeria Panzetta, Cristina D’Aniello

## Abstract

Pancreatic ductal adenocarcinoma (PDAC) develops within a desmoplastic and stiffened microenvironment that critically shapes tumor progression and therapeutic resistance, yet these features are not reproduced by conventional rigid plastic culture systems. Here, we leverage a tuneable bioengineered platform that mimics stromal stiffening to investigate how mechanical cues regulate PDAC cell behaviour and to identify pharmacological strategies that counteract stiffness-driven malignancy. We show that increasing matrix stiffness promotes key hallmarks of PDAC aggressiveness, including enhanced cell spreading, focal adhesions maturation, and cytoskeletal tension. Notably, we identify the glucocorticoid budesonide as a selective suppressor of stiffness-induced malignant phenotypes. Transcriptomic profiling reveals that budesonide counteracts stiffness-associated gene programs, prominently affecting pathways governing cytoskeletal dynamics, nuclear envelope organization, and YAP nucleocytoplasmic transport. Consistently, budesonide reduced force transmission to the nucleus, restoring nuclear wrinkling and constraining nuclear size and shape. These effects are mediated through both glucocorticoid receptor-dependent and -independent mechanisms, revealing a previously unrecognized mode of action. Together, our findings establish mechanical context as a critical determinant of PDAC vulnerability and identify budesonide as a candidate for therapeutic repurposing to target stiffness–driven cancer progression.

## INTRODUCTION

Pancreatic ductal adenocarcinoma (PDAC) is a deadly disease that is predicted to become the second leading cause of cancer–related death by 2030^1^. The dismal prognosis of pancreatic cancer is primarily caused by the fact that 90% of tumours are diagnosed at an advanced stage, with systemic metastases in over 50% of cases, compounded by the absence of effective screening methods^2^. The hallmark of PDAC is the profoundly altered tumour mechanical microenvironment (TMME)^3^, characterized by extensive collagen deposition and extracellular matrix (ECM) remodeling^4^. The desmoplastic ECM, enriched in fibrillar collagen and glycosaminoglycans^5^, evolves into a highly organized and stiff structure^5–7^, fostering physical and biochemical cell–cell and cell–matrix interactions that drive tumour progression^5,7,8^ and therapy resistance^6,9–12^. Cells sense alterations in the mechanical properties of the TMME and undergo coordinated structural changes^13^ from focal adhesions that mediate direct interactions with the ECM, to the cytoskeleton and its force–generating machinery, up to the nucleus, where mechanical inputs can regulate transcriptional activity^14–16^. As a result, these mechanical forces are transduced into intracellular signalling programs via mechano–transduction pathways, among which the YAP/TAZ complex acts as a key transcriptional effector driving proliferation, survival and therapy resistance^17–21^.

Building on these insights, increasing efforts have been focused on the development of therapeutic approaches aimed at counteracting stiffness–induced aggressive traits in cancer cells. However, molecules able to avoid the influence of the TMME on cancer cells are so far not available^22^, in part due to the lack of model systems that accurately mimic the TMME. In this context, the development of bioengineered platforms that better mimic the impact of the stiff TMME on cancer cells is becoming an increasing field of interest^23,24^.

Our work presents an experimental approach to model PDAC aggressive features through mechanobiology principles^18,20^, using bioengineered platforms that recreate TMME conditions with controlled stiffness. Beyond providing a more physiologically relevant model, this platform also serves as a predictive tool for therapeutic testing. Among candidate compounds, the FDA–approved glucocorticoid budesonide, widely used in the treatment of asthma^25^ and inflammatory diseases^26^, has recently emerged as a context–dependent regulator of PDAC cell behavior^27^, suggesting a previously unidentified antitumor activity beyond its well–established role as an anti–inflammatory glucocorticoid^28^.

Here, we find that budesonide antagonizes the acquisition of the stiffness–driven malignant traits in PDAC by remodelling cytoskeletal organization and modulating nuclear architecture, signalling pathways, and transcriptional programs. This coordinated regulation suppresses aggressive PDAC cell features, including migration and proliferation, and identifies budesonide as a candidate therapeutic strategy for PDAC treatment.

## RESULTS

### Stiffer substrates promote aggressive behaviour in pancreatic cancer cells

To interrogate the mechanosensory capacity of pancreatic ductal adenocarcinoma (PDAC) cells, we engineered polyacrylamide (PAAm) hydrogels spanning a physiologically relevant stiffness range (Young’s moduli: 0.2, 10, and 27 kPa; Supplementary Fig. 1A). These substrates recapitulate distinct mechanical states of the pancreatic microenvironment, with soft matrices (0.2 kPa) reflecting physiological or sub–physiological tissue compliance, and intermediate (10 kPa) to stiff (27 kPa) matrices modelling progressive stromal remodelling associated with disease progression^29^. While prior studies have examined selected pancreatic cell responses to matrix stiffness, a comprehensive view of the coordinated biological, mechanical, and molecular adaptations elicited across defined mechanical contexts is still lacking. To address this gap, we performed an integrated characterization of PDAC cell responses to tuneable substrate stiffness.

PANC-1 cells were first seeded on gelatin–coated substrates (1.5 × 10⁴ cells/cm²) and, after 72 h, detached and replated at low density (2.5 × 10³ cells/cm²) onto PAAm hydrogels of defined stiffness. Cells were analysed 48 h post–seeding (Supplementary Fig. 1B). To determine how substrate mechanics regulates cell–matrix interactions, we quantified focal adhesion organization and cell spreading by immunofluorescence imaging of paxillin and vimentin/F–actin, respectively. PANC-1 cells cultured on soft substrates (0.2 kPa) displayed sparse and poorly developed focal adhesions, indicative of weak cell–matrix coupling (Fig. 1A, B). In contrast, increasing substrate stiffness promoted the formation of elongated, well–defined adhesion complexes, accompanied by a progressive increase in focal adhesion area (Fig. 1A, B), consistent with strengthened adhesive engagement. Accordingly, cell spreading progressively increased with substrate stiffness, as reflected by a marked increase in projected cell area (Supplementary Fig. 1C). This was accompanied by the emergence of prominent actin stress fibres and a reinforced vimentin network, supporting enhanced cytoskeletal tension and mechanotransductive coupling with the ECM (Supplementary Fig. 1C). The observed cytoskeletal remodelling was paralleled by changes in cellular mechanics, as quantified by atomic force microscopy (AFM)–based force spectroscopy (Fig. 1C). PANC-1 cells progressively increased their stiffness in response to substrate rigidity (median: 0.31 kPa at 0.2 kPa; 0.62 kPa at 10 kPa; 1.01 kPa at 27 kPa), indicating adaptive mechanical tuning to the extracellular environment (Fig. 1C).

**Figure 1.**
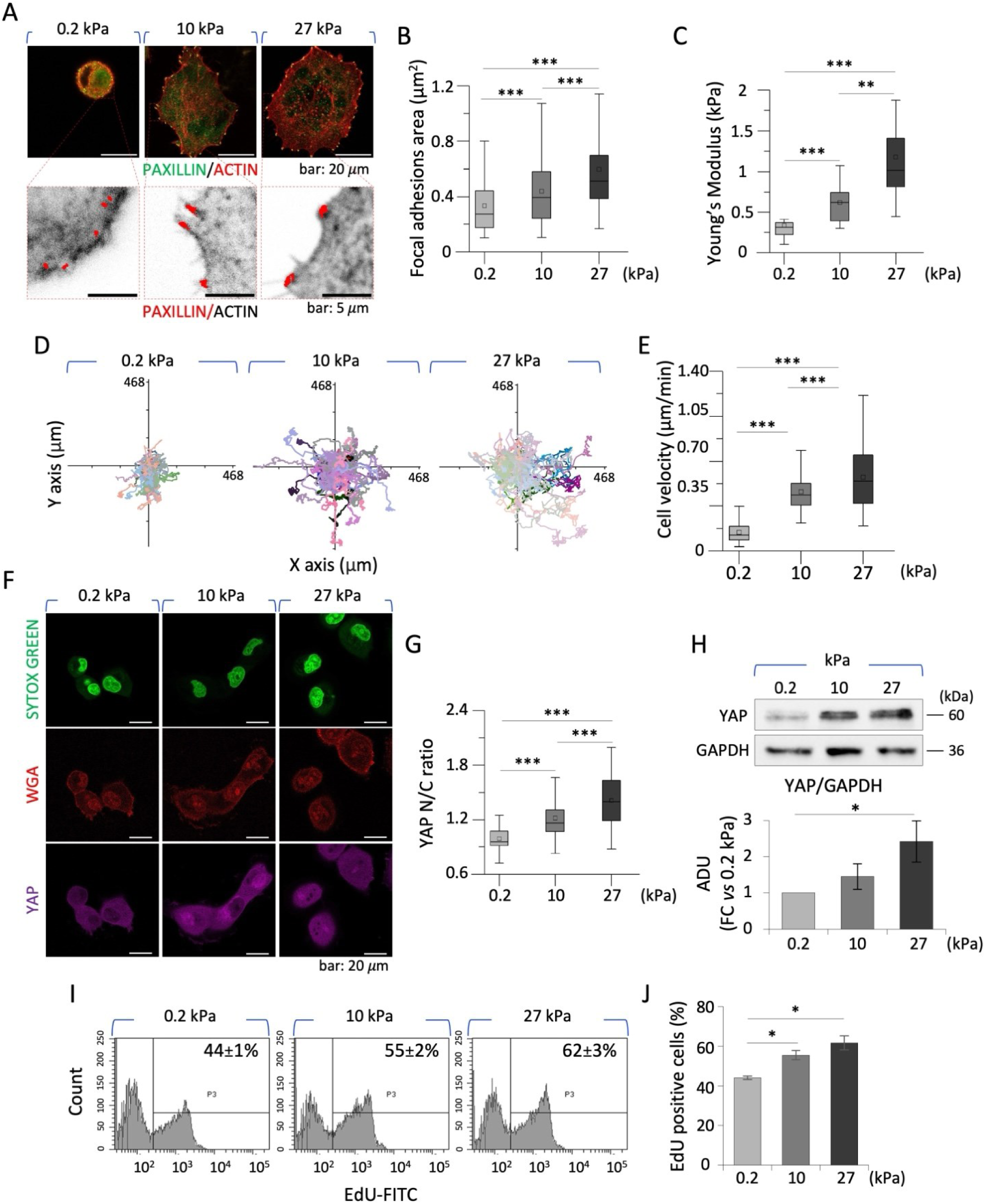
Pancreatic cancer cells sense increasing substrate stiffness. **A-B** Representative confocal images (A) and quantification (B) of PANC-1 cells, seeded on the bioengineered platforms at 0.2, 10 and 27 kPa, showing paxillin (green) and F-actin filaments (red) staining with focal adhesions highlighted in yellow (*upper*). The corresponding zoomed inverted contrast views with focal adhesions’ ROIs selected from paxillin (red) and F-actin (black) (*lower*) are shown underneath, with the corresponding focal adhesion area analyses plot (B, n=30). **C** Young’s modulus evaluation, using AFM in force spectroscopy mode, of PANC-1 cells (n=15) two days after seeding on the bioengineered platforms. **D-E** Plotted single-cell trajectories of PANC-1 cells (D) centred in the axes’ origin and single-cell velocity analyses in time lapse microscopy (E, n=100). **F**-**G** Representative confocal images of YAP (purple) and WGA-rhodamine (red) staining (F), with the corresponding YAP N/C ratio intensity analyses (G, n=30), in PANC-1 cells. Nuclei were stained with SYTOX green (green). **H** Representative western blot analysis (*upper*) and densitometric quantification (FC vs 0.2; *lower*) of YAP in PANC-1 cells at different mechanical conditions. GAPDH was used as a loading control. **I-J** Representative FACS plot of EdU incorporation (I) and Box plot (J) showing percentage of EdU-positive cells cultured on 0.2, 10, and 27 kPa substrates. Western blot data are shown as mean ± SEM (N=3). All other quantitative data are presented as box plots (N=3-5). Significant statistical differences are reported: * p < 0.05; ** p < 0.01; *** p < 0.001. The complete statistical analysis executed on shown datasets can be found in Supplementary Table 5.

Stiffness–dependent reinforcement of focal adhesions and cytoskeletal organization are key determinants of traction force generation and cell motility^30,31^. Thus, we next investigated how substrate mechanics regulates PANC-1 cell migration. Time–lapse microscopy combined with single–cell tracking over 48 h revealed that increasing substrate stiffness enhanced migratory behaviour. Trajectories aligned to a common origin qualitatively demonstrated an expansion of the explored area with increasing stiffness (Fig. 1D). Of note, cell velocity exhibited a non–linear increase (median: 0.12 µm min⁻¹ at 0.2 kPa; 0.43 µm min⁻¹ at 10 kPa; 0.52 µm min⁻¹ at 27 kPa; Fig. 1E), paralleled by higher mean square displacement (MSD), indicating greater spatial exploration (Supplementary Fig. 1E). In contrast, the slope of the MSD curves appeared largely independent of substrate stiffness, suggesting that migration remained predominantly random, consistent with the isotropic mechanical and biochemical properties of the substrates, which do not provide cues for directional movement.

Given the close interplay between cytoskeletal tension and mechanosensitive transcriptional programs^32,33^, we next investigated the subcellular localization of the key mechanotransducers YAP/TAZ^17,21^. Immunofluorescence analysis revealed a stiffness–dependent increase in YAP nuclear accumulation, as reflected by a non–linear rise in the nuclear–to–cytoplasmic (N/C) ratio (median: 0.96 at 0.2 kPa; 1.16 at 10 kPa; 1.40 at 27 kPa; Fig. 1F, G). These findings were further supported by a progressive increase in total YAP protein levels with substrate stiffness (Fig. 1H), as expected^34–37^. Then, we assessed how cell proliferation was modulated by increasing substrate stiffness. EdU incorporation assays revealed a significant, stiffness–dependent increase in DNA synthesis, with a higher fraction of EdU–positive cells on stiffer substrates (44 ± 1% at 0.2 kPa; 55 ± 2% at 10 kPa; 62 ± 3% at 27 kPa; Fig. 1I, J). Consistent with these findings, time–lapse microscopy showed a progressive increase in proliferation rates with increasing stiffness (median: 0.20, 0.39, and 0.50 doublings/day at 0.2, 10, and 27 kPa, respectively; Supplementary Fig. 1F), suggesting that a mechanically rigid microenvironment promotes both the mechanical adaptation and proliferative expansion of PANC-1 cells.

Collectively, these findings advance current understanding by providing a comprehensive, multi–scale characterization of pancreatic cancer cell mechanoadaptation, highlighting how matrix stiffness modulates key cellular behaviours associated with malignancy.

### Budesonide treatment counteracts the stiffness–mediated effects in pancreatic cancer cells

We next asked whether the tuneable bioengineered platforms could be exploited to identify compounds that counteract stiffness–driven PDAC phenotypes. We focused on the glucocorticoid budesonide, recently proposed as a candidate for drug repurposing in PDAC^26,27^. Notably, budesonide exerts marked context–dependent effects, suppressing PDAC cell proliferation only in 3D models, while restraining mesenchymal and invasive traits in conventional 2D cultures^26,27^. However, the mechanisms underlying these responses remain incompletely understood. Given the strong influence of the microenvironment on budesonide activity^27^, we reasoned that bioengineered platforms recapitulating the physical and biochemical properties of the tumour microenvironment would provide a more relevant framework than standard rigid plastic substrates.

PANC-1 cells were therefore seeded as described (Supplementary Fig. 2A), treated with budesonide or dexamethasone (20 µM), or left untreated (NT) as control. After three days in culture, cells (2.5×10^3^ cells/cm^2^) in presence or absence of budesonide or dexamethasone were replated onto stiff substrates (27 kPa) to model the pathological PDAC microenvironment (Supplementary Fig. 2A). NT cells cultured on soft substrates (0.2 kPa) served as a reference. Immunofluorescence analysis of vimentin, F–actin and paxillin staining revealed that budesonide markedly attenuated stiffness–induced cytoskeletal remodelling, reducing both focal adhesion area and cell spreading (Fig. 2A, B and Supplementary Fig. 2B, C). This was accompanied by the emergence of a cortical actin ring and a concomitant loss of internal stress fibres (Supplementary Fig. 2B, C). In contrast, dexamethasone–treated cells resembled NT cells on stiff substrates, with a comparable spreading area, but only a modest reduction in focal adhesions size compared to budesonide–treated cells (Fig. 2A, B and Supplementary Fig. 2B, C), thus suggesting a partial GR–independent mechanism of budesonide action or different receptor affinities. To address this issue and assess the involvement of the GR, we generated PANC-1 cells silenced for *NR3C1*, the gene encoding the GR^38^. Briefly, PANC-1 cells were infected with lentiviruses carrying either a short hairpin (sh) against the *NR3C1* gene, or an empty vector as control. Western blot analysis confirmed an efficient downregulation of *NR3C1* protein, with a ∼75% reduction in PANC-1 cells silenced for *NR3C1* (*^shNR3C1^*PANC-1) compared to control cells (*^shEmpty^*PANC-1; Supplementary Fig. 3A). We then investigated whether GR knockdown affected the response to budesonide by culturing *^shEmpty^*PANC-1 and *^shNR3C1^*PANC-1 cells with or without budesonide (20 µM). After three days in culture, cells were dissociated and replated (2.5×10^3^ cells/cm^2^) on stiff substrates and cultured for additional two days (Supplementary Fig. 3B). We first assessed the effect of GR knockdown on cell adhesion. Focal adhesions contact area measurements and cell spreading area showed that budesonide strongly affected both parameters in *^shEmpty^*PANC-1 cells (Supplementary Fig. 3C-E). Interestingly, GR silencing reduced focal adhesion contact and spreading area, while further exposure to budesonide led only to a slight reduction of focal adhesion, without significantly affecting overall cell spreading (Supplementary Fig. 3C-E), suggesting that GR signalling may contribute to cytoskeletal organization.

**Figure 2.**
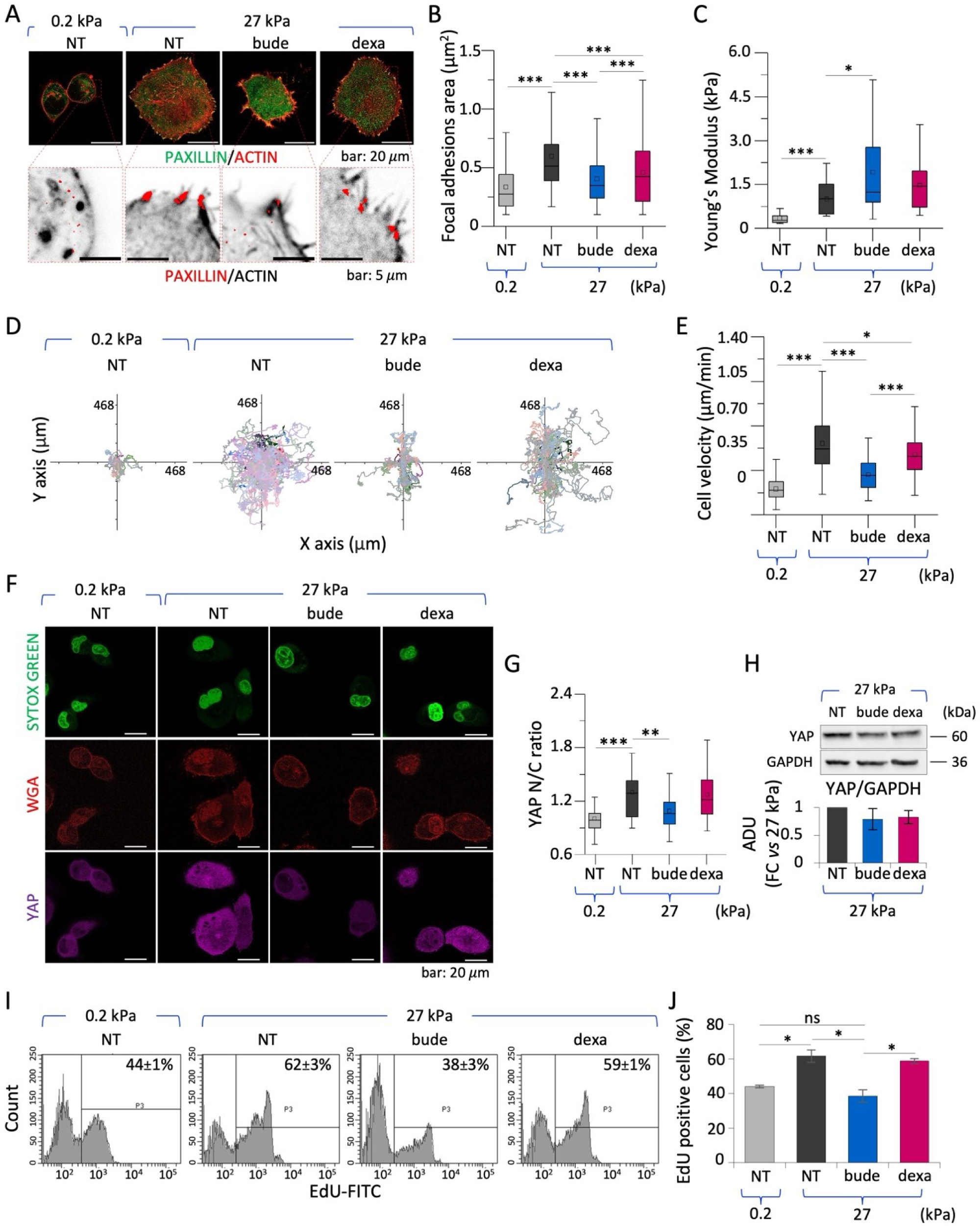
Budesonide treatment counteracts the stiffness-mediated effects in PANC-1 cells. **A-B** Representative confocal images (A) and quantification (B) of PANC-1 cells plated on 0.2 and 27 kPa substrates and treated ± budesonide or dexamethasone (20 μM), showing paxillin (green) and F-actin filaments (red) staining with focal adhesions highlighted in yellow (*upper*) and the corresponding zoomed inverted contrast views with focal adhesions’ ROIs selected from paxillin (red) and F-actin (black) (*lower*). The quantitative dataset from focal adhesions area analyses is shown in B (n=30). **C** Young’s modulus evaluation of PANC-1 cells after treatment, as above, using AFM in force spectroscopy mode two days after seeding (n=15). **D-E** Plotted single-cell trajectories centred in axes origin of treated PANC-1 cells (*E*), and single-cell velocity analyses in time lapse microscopy (D, n=100). **F-G** Representative confocal images (F) of YAP (purple) and WGA-rhodamine (red) staining, with the corresponding YAP N/C ratio intensity analyses (G, n=50), in PANC-1 cells after treatment. Nuclei were stained with SYTOX green (*green*). **H** Representative western blot analysis (*upper*) and densitometric quantification (FC vs 27 kPa NT; *lower*) of YAP in treated PANC-1 cells. GAPDH was used as a loading control. **I-J** Representative FACS plot of EdU incorporation (I) and box plot (J) showing percentage of EdU-positive cells cultured on 0.2 and 27 kPa substrates ± budesonide or dexamethasone (20 μM). Western blot data are shown as mean ± SEM (n=3). All other quantitative data are presented as box plots (N=3-5). Significant statistical differences are reported: * p < 0.05; ** p < 0.01; *** p < 0.001. The complete statistical analysis executed on shown datasets can be found in Supplementary Table 7.

Consistent with these structural changes, AFM measurements showed that budesonide altered cellular mechanical properties (Fig. 2C). The observed increase in apparent elastic modulus is largely attributable to cortical actin reorganization rather than enhanced internal cytoskeletal tension typically induced by stiffness. By contrast, dexamethasone did not significantly alter PANC-1 mechanics, supporting the specificity of budesonide.

Given the observed changes in adhesion dynamics, we further examined cell motility. Single–cell tracking revealed that budesonide significantly reduced migration on stiff substrates, both in terms of velocity and the extent of spatial exploration (Fig. 2D, E and Supplementary Fig. 2D). In contrast, dexamethasone induced only a modest, albeit significant, reduction of cell velocity (Fig. 2D, E). Consistently, MSDs of PANC-1 cells exposed to dexamethasone largely overlapped with those of NT cells at the same stiffness (Supplementary Fig. 2D). This convergence suggests that dexamethasone–treated cells explore a larger effective area over time, comparable to NT cells on stiff substrates. By contrast, budesonide markedly reduced cell displacement, reaching levels close to those observed on soft substrates, consistent with a more spatially confined migration pattern (Supplementary Fig. 2D). Despite these differences in displacement amplitude, migration remained predominantly random across all conditions, consistent with the absence of directional cues. We next investigated the impact of GR knockdown on cell motility by time–lapse analysis. As expected, budesonide was able to prevent the stiffness–dependent increase of cell motion velocity and area coverage in *^shEmpty^*PANC-1 cells compared to NT (Supplementary Fig. 3F, G). Interestingly, we observed a decreased average cell velocity in *^shNR3C1^*PANC-1 cells, not further reduced by budesonide supplementation (Supplementary Fig. 3F, G). These findings support a previously unrecognized role of the GR in regulating pancreatic cancer cell motility. However, they do not allow a clear distinction between GR–dependent and GR–independent mechanisms underlying budesonide–induced cytoskeletal remodelling and migration.

We next assessed the effect of budesonide on YAP nuclear translocation. As expected, YAP was predominantly nuclear in NT cells on stiff substrates. Budesonide treatment promoted YAP cytoplasmic retention, reducing the N/C ratio to levels approaching those observed on soft substrates (Fig. 2F, G), without altering total YAP protein levels (Fig. 2H). These findings indicate that budesonide primarily modulates YAP localization rather than overall abundance. No significant effect on YAP nuclear translocation was observed with dexamethasone. Finally, we assessed the impact of budesonide on cell proliferation by EdU incorporation assay. Budesonide counteracted the stiffness–induced increase in EdU incorporation, restoring proliferation rates to levels comparable to those observed on soft substrates (Fig. 2I, J). Time–lapse analysis confirmed that budesonide prevented the stiffness–driven expansion of PANC-1 cells, whereas dexamethasone exerted only a partial effect (Supplementary Fig. 2E). We thus compared the growth rate of *^shEmpty^*PANC-1 and *^shNR3C1^*PANC-1 cells using time–lapse imaging. Both cell lines exhibited similar growth rates, indicating that knockdown of GR does not impact PANC-1 cell growth (Supplementary Fig. 3H). Budesonide significantly counteracted the stiffness–dependent proliferation induction in both *^shEmpty^*PANC-1 and *^shNR3C1^*PANC-1 cells under these conditions (Supplementary Fig. 3H), supporting a potential contribution of a GR–independent mechanism in budesonide activity.

Collectively our data reveal a previously unappreciated role for the GR in regulating cytoskeletal organization and migration in pancreatic cancer cells and led us to hypothesize that budesonide counteracts stiffness–induced proliferation through a GR–independent mechanism. Furthermore, they establish budesonide as a suppressor of stiffness–driven malignancy and underscore the need for physiologically relevant mechanical models to uncover otherwise hidden drug responses.

### Budesonide treatment counteracts the stiffness–mediated effects in patient–derived PDAC cells

To validate our findings in clinically relevant models, we extended our analysis to patient–derived PDAC cell lines (PDAC#253 and PDAC#354 cells^39,40^), which more faithfully recapitulate disease–specific traits. PDAC#253 and PDAC#354 cells were seeded on gelatin–coated substrates (1.5 × 10⁴ cells/cm^2^) and treated 24 h post–plating with budesonide or dexamethasone (20 µM) or left untreated (NT). After 72 h, cells were dissociated and replated at low density (2.5 × 10³ cells/cm^2^) onto stiff substrates and cultured for additional 48 h. NT cells seeded on soft substrates (0.2 kPa) served as controls (Fig. 3A). Consistent with findings in PANC-1 cells, patient–derived PDAC cells exhibited pronounced stiffness–dependent changes in morphology and cytoskeletal organization (Fig. 3B, C and Supplementary Fig. 4A-D). On stiff substrates, NT cells appeared more elongated and with a spread morphology compared to those seeded on the soft substrate. In contrast, budesonide–treated cells adopted a rounded shape, with reduced focal adhesion area and a disorganized cytoskeleton, resembling cells cultured on soft substrates (Fig. 3B, C and Supplementary Fig. 4A-D). These changes were accompanied by the emergence of aberrant cortical actin structures (Supplementary Fig. 4A- D). In contrast, dexamethasone had minimal impact on focal adhesion architecture and cytoskeletal organization, with cells largely retaining a stiff–like phenotype (Fig. 3B, C and Supplementary Fig. 4A-D).

**Figure 3.**
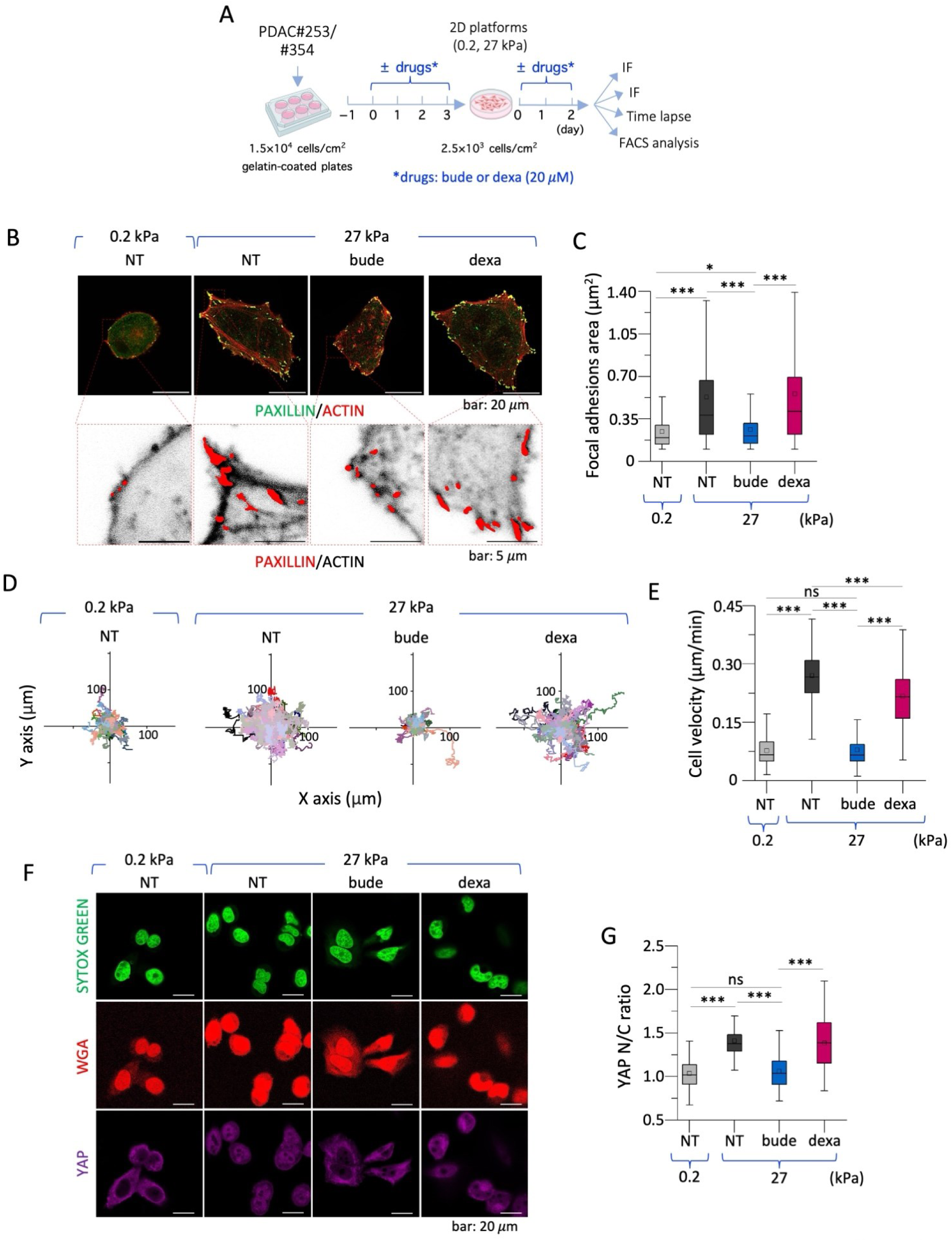
Budesonide treatment counteracts the stiffness-mediated effects in patient-derived PDAC cells. **A** Schematic representation of the experimental procedure. PDAC#253 cells were first plated on gelatin-coated plates at 1.5×10^4^ cells/cm^2^ at day –1. On day 0 cells were treated ± budesonide (20 µM) or dexamethasone (20 µM) for three days. Cells were then detached with trypsin and plated at 2.5×10^3^ cells/cm^2^ on the 27 kPa substrate for two days. Control NT cells were also plated on the 0.2 kPa substrate. **B-C** Representative confocal images (B) of PDAC#253 cells treated as described in A, showing paxillin (green) and F-actin filaments (red) staining with focal adhesions highlighted in yellow (*upper*). The corresponding zoomed inverted contrast views with focal adhesions’ ROIs selected from paxillin (red) and F-actin (black) are shown underneath (*lower*). Focal adhesions’ area analyses highlight dimensional dependency on substrate stiffness and drug treatment (C, n=20). **D-E** Plotted single-cell trajectories centred in axes origin (D) and single-cell velocity analyses in time lapse microscopy (E, n=125) of PDAC#253 cells upon treatment. **F-G** Representative confocal images (F) of YAP (purple) and WGA-Rhodamine (red) staining, with the corresponding YAP N/C ratio intensity analyses (G, n=60), of treated PDAC#253 cells. Nuclei were stained with SYTOX green (*green*). Quantitative data are presented as box plots (N=3). Significant statistical differences are reported: * p < 0.05; ** p < 0.01; *** p < 0.001. The complete statistical analysis executed on shown datasets can be found in Supplementary Table 10.

We next assessed proliferative capacity by time–lapse imaging. Budesonide consistently counteracted the stiffness–induced increase in proliferation in both PDAC#253 and PDAC#354 cells, restoring growth rates to levels comparable to those observed on soft substrates (Supplementary Fig. 4E, F). By contrast, dexamethasone showed negligible or modest effects, depending on the cell line. Similarly, single–cell tracking revealed that budesonide effectively suppressed the stiffness–driven increase in cell motility across both patient–derived cells (Fig. 3D, E and Supplementary Fig. 4G, H), whereas dexamethasone produced only partial reductions with consistent effect across cell lines.

Finally, we examined YAP mechanotransduction. As expected, YAP nuclear localization increased on stiff substrates, confirming a conserved mechanosensitive response in patient–derived PDAC cells (Fig. 3F, G and Supplementary Fig. 4I, J). Budesonide significantly reduced YAP nuclear translocation in both PDAC cells, restoring N/C ratios towards soft–like conditions. In contrast, dexamethasone produced minimal effects overall, with responses varying across patient–derived cell lines (Fig. 3F, G and Supplementary Fig. 4I, J).

Collectively, these findings demonstrate that budesonide consistently suppresses stiffness–driven malignant phenotypes across different patient–derived PDAC models, reinforcing its role as a potent and selective inhibitor of PDAC mechanoadaptation.

### Budesonide prevents the stiffness–driven transcriptional changes and impacts nucleoskeletal organization of pancreatic cancer cells

To gain further mechanistic insight into the effects of budesonide on stiffness–driven PDAC phenotypes, we performed RNA–seq profiling of PANC-1 cells cultured on soft and stiff substrates, as well as budesonide–treated cells maintained on stiff substrates (Fig. 4A). Principal component analysis revealed clear segregation of the three conditions, indicating distinct transcriptional states (Fig. 4B). Comparison of PANC-1 cells on stiff versus soft substrates identified 2073 differentially expressed genes (Supplementary Fig. 5A), reflecting transcriptional adaptation to increased matrix stiffness. David gene ontology and pathway enrichment (padj≤0.05) analyses of deregulated genes highlighted processes linked to regulation of metabolic process, cytoskeletal organization, epithelial to mesenchymal transition, regulation of cell adhesion mediated by integrin, migration, and intracellular signalling, as well as enrichment of Hippo signalling and cell cycle pathways (Fig. 4C, D; Supplementary Fig. 5B, C; Supplementary Table 1), consistent with the phenotypic changes observed.

**Figure 4.**
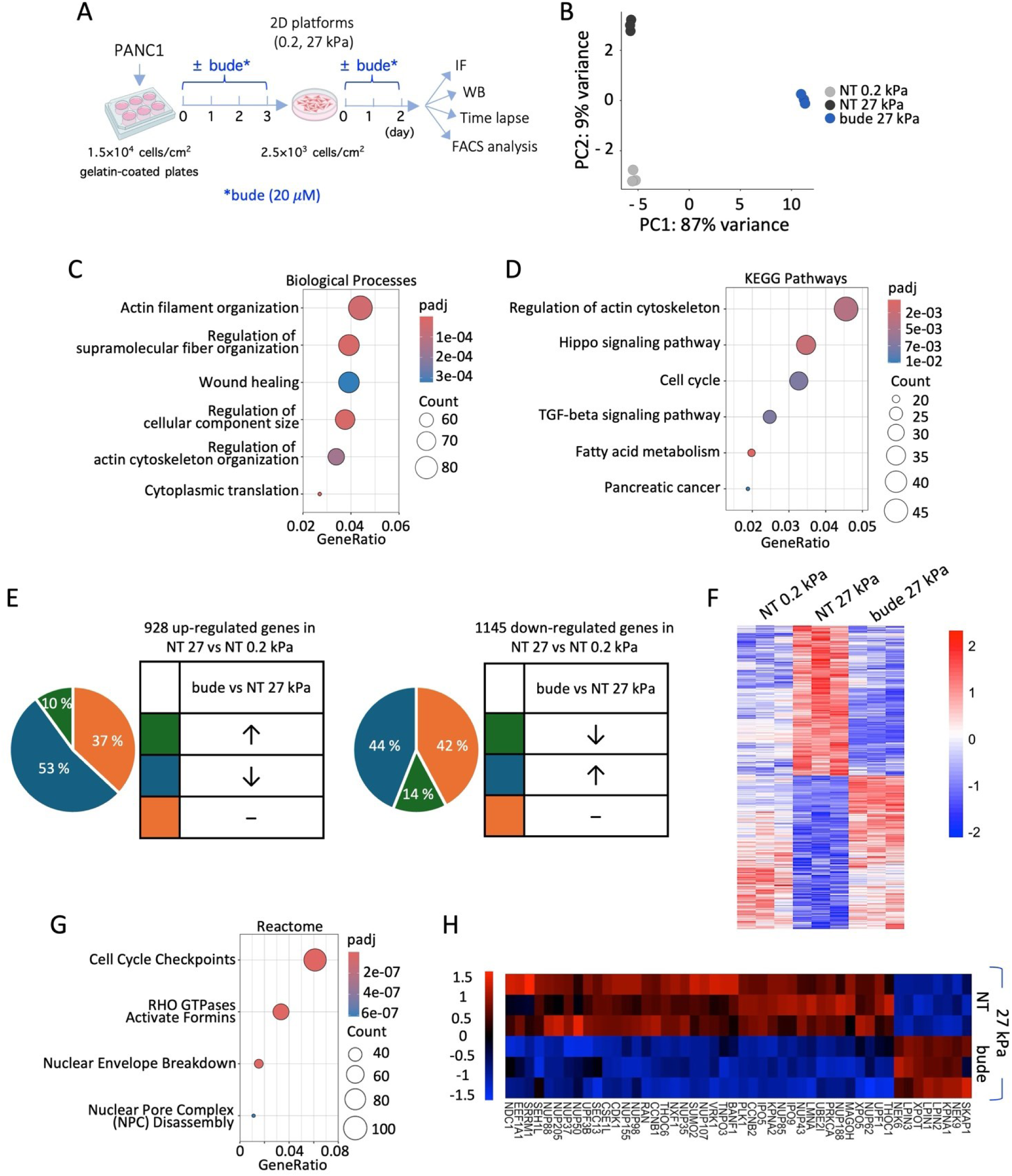
Transcriptome profiling of PANC-1 cells plated on the bioengineered platforms and treated with budesonide. **A** Schematic representation of the experimental procedure. PANC-1 cells were first plated on gelatin-coated plates at 1.5×10^4^ cells/cm^2^, ± budesonide (20 µM). After three days in culture, cells were detached with trypsin and plated at 2.5×10^3^ cells/cm^2^ on the stiffer 27 kPa substrate for additional two days, ± budesonide (20 µM). Control NT cells were also plated on the 0.2 kPa substrate. **B** Principal component analysis (PCA) of the transcriptomic profile of PANC-1 cells in the determined mechanical and treatment conditions. **C-D** Enrichment data analysis of differentially expressed genes for biological processes (C) and KEGG Pathways (D) in the comparison between NT cells plated on the 27 *versus* the 0.2 kPa substrates. **E** Pie graphs showing the percentage of the deregulated genes that are inversely regulated by stiffness and budesonide. **F** Heatmap of 997 differentially expressed genes of PANC-1 cells cultured and treated as described in A. **G** Enrichment data analyses of differentially expressed genes for Reactome specifically modulated by budesonide (∼3000 genes) in PANC-1 cells seeded on stiff substrates. **H** Heatmap of the deregulated genes related to nuclear envelope breakdown, nuclear pore complex disassembly, nucleocytoplasmic transport in the between budesonide-treated and untreated (NT) cells, seeded on the 27 kPa substrates.

We next compared the transcriptomic profiles of PANC-1 cells cultured on stiff substrates following budesonide treatment or vehicle control and identified 4271 differentially expressed genes (Supplementary Fig. 5A). Of note, cross–comparison with stiffness–regulated genes (2073; Supplementary Fig. 5A) revealed that approximately 50% of the stiffness–deregulated genes were inversely modulated by budesonide (Fig. 4E, F; Supplementary Table 2), supporting its ability to counteract stiffness–driven programs associated with malignant phenotypes.

To further dissect budesonide–specific effects, we analysed genes uniquely altered by budesonide (∼3000 genes). Enrichment analyses using Reactome (Fig. 4G), BP (Supplementary Fig. 5D) and KEGG Pathways (Supplementary Fig. 5E) revealed modulation of pathways related to cell cycle regulation, consistent with the anti–proliferative effects of budesonide (Supplementary Table 1). Unexpectedly, we also identified a distinct enrichment in pathways related to nuclear envelope organization, nuclear pore complex dynamics, and nucleocytoplasmic transport (Fig. 4G, H and Supplementary Fig. 5D, E; Supplementary Table 2), pointing to a previously unappreciated impact of budesonide on nuclear architecture.

To validate these observations, and establish their relevance in patient–derived models, we examined nuclear lamina organization by immunofluorescence analysis of lamin A/C and lamin B1 in both PANC-1 and PDAC#253 cells. In both models, budesonide reduced lamin A/C levels compared to NT cells cultured on stiff substrates, while lamin B1 showed a cell line–dependent response (Fig. 5A-F), suggesting mechanosensitive–dependent regulation of these nucleoskeleton components^41^.

**Figure 5.**
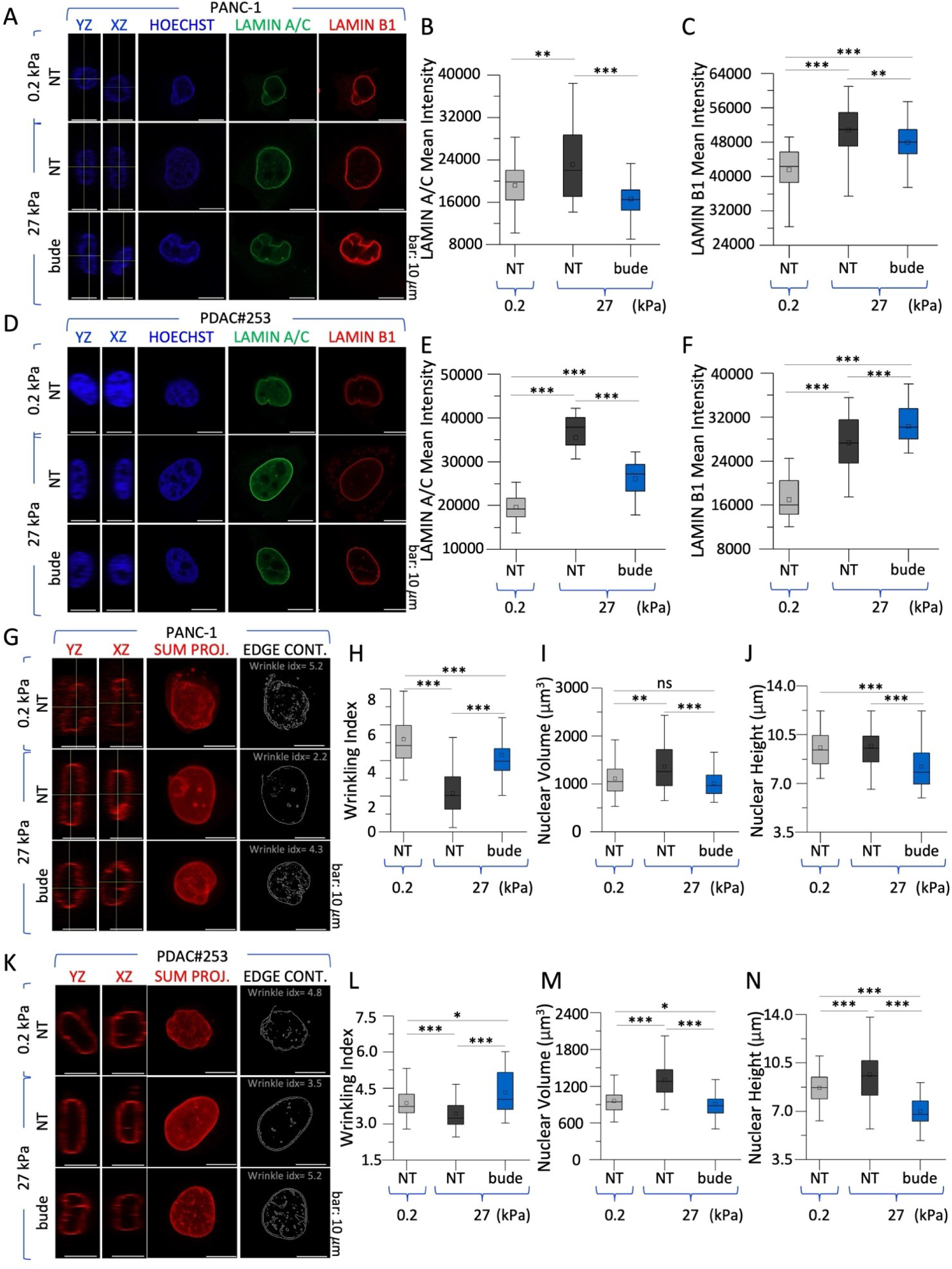
Budesonide modifies the nuclear architecture in PANC-1 and patient-derived cells. **A-F** Representative confocal images of lamin A/C (green) and lamin B1 (red) in PANC-1 (A) and PDAC#253 (D) cells plated as described in Supplementary Fig. 2A. Nuclei were counterstained with HOECHST (blue). The orthogonal projections in the YZ and XZ planes of representative nuclei are shown (A, D), alongside the quantifications of lamin A/C (B, E) and lamin B1 (C, F) mean fluorescence intensities. **G-I, K-M** Representative confocal images of the sum projection, alongside the orthogonal projections in the YZ and XZ planes for lamin B1 (red) of PANC-1 (G) and PDAC#253 cells (K). The edge contour representation extracted through Sobel-based edge filtering with the wrinkling index corresponding to the representative image (G, K), the quantification of the wrinkling indexes (H, L) and the nuclear volume (I, M) are shown. **J, N** Quantitative morphometry of PANC-1 cells nuclei, obtained by measurement of the bounding box encompassing the nuclear signal extracted from HOECHST staining, shown as box plots. Scale bar: 10 µm. Quantitative data are presented as box plots (n=35, N=3). Significant statistical differences are reported: * p < 0.05; ** p < 0.01; *** p < 0.001. The complete statistical analysis executed on shown datasets can be found in Supplementary Table 13 and 14.

To further explore this phenotype, we next assessed nuclear membrane topology by quantifying nuclear membrane wrinkling. Cells on soft substrates exhibited higher wrinkling indices than those on stiff matrices (Fig. 5G, H, K, L), as expected^33,41,42^. Notably, budesonide significantly increased nuclear wrinkling index towards soft–like levels in both PANC-1 and PDAC#253 cells (Fig. 5G, H, K, L), consistent with reduced transmission of cytoskeleton mechanical force to the nucleus.

Alterations in the enveloping laminar structure around the nucleus significantly impacts nuclear size and shape^33,43,44^. Consistent with these structural changes, budesonide significantly reduced nuclear volume across the different models, restoring values comparable to those observed under soft conditions (Fig. 5I, M). To further dissect these changes, we quantified nuclear height and the major and minor axes of the nuclear midplane. Budesonide treatment significantly reduced nuclear height and major axis in both PANC-1 and PDAC#253 cells, while a decrease in the minor axis was observed only in PDAC#253 cells (Fig. 5J, N and Supplementary Fig. 6A-B), which may reflect a cell–line specific effect.

We next asked whether the impact of budesonide on nuclear organization involves the GR. Immunofluorescence analysis of nuclear lamina components showed that, as expected, budesonide reduced both lamin A/C and lamin B1 expression in *^shEmpty^*PANC-1 cells (Supplementary Fig. 6C-E). GR knockdown itself reduced the levels of both laminae and budesonide did not further reduce their expression in *^shNR3C1^*PANC-1 cells (Supplementary Fig. 6C-E). Similarly, GR loss increased nuclear surface wrinkling, with no additional effect upon budesonide treatment, suggesting a GR involvement in this process (Supplementary Fig. 6F, G). In contrast, GR downregulation did not affect nuclear volume, major and the minor axes and the nuclear height. Budesonide, however, significantly reduced these parameters in both *^shEmpty^*PANC-1 and *^shNR3C1^*PANC-1 cells, suggesting that the effects of budesonide on nuclear size and shape are, at least in part, GR–independent (Supplementary Fig. 6H-K). Notably, despite recapitulating several soft–like features, including reduced adhesion, reduced migration and proliferation, and altered lamina organization, GR knockdown did not affect nuclear volume. This suggests that global nuclear geometry is not solely dictated by lamina composition or cytoskeletal tension but instead requires coordinated changes across multiple determinants of nuclear architecture, with nuclear volume representing a higher–order emergent property that is not readily altered by isolated changes in cytoskeletal tension or lamina organization.

Collectively, these findings provide unprecedented evidence that budesonide reverses stiffness–driven transcriptional programs associated with malignant phenotypes, and remodels nuclear architecture in PDAC cells, thereby attenuating mechanotransduction at the nuclear level.

## DISCUSSION

Matrix stiffening is a defining feature of PDAC and a key driver of malignant cell behaviour through activation of mechanotransductive pathways. Here, using tuneable bioengineered platforms that recapitulate physiologically relevant stiffness ranges, we identify the FDA–approved glucocorticoid budesonide as a previously unrecognized modulator of PDAC mechanobiology. Budesonide effectively counteracts stiffness–driven phenotypes, including enhanced cell spreading, focal adhesion maturation, cytoskeletal organization, migration and proliferation^7,45–48^. These findings demonstrate that pharmacological targeting of mechanoadaptive programs can be achieved using an already clinically available compound and highlight the importance of incorporating mechanical cues into preclinical models.

While individual aspects of stiffness–induced responses, such as YAP activation^17–21^, cytoskeletal remodelling^6,7,13,14,47^ and increased proliferation^45,49^, have been described in various tumour contexts^18,20,48,50^, a comprehensive and integrated characterization of these processes in PDAC has remained limited. By decoupling matrix stiffness from other microenvironmental variables, our tuneable platforms enabled a multi–scale analysis linking extracellular mechanics to cytoskeletal organization, nuclear architecture and transcriptional regulation. Within this framework, matrix stiffening induces a coordinated phenotypic program consistent with the establishment of a self–reinforcing mechanotransductive circuitry^8,16,32,46,51–53^.

A central insight of this study is that budesonide disrupts stiffness–induced mechanotransductive programs. Our data indicate that this occurs early at the level of cytoskeletal organization, impairing force transmission toward the nucleus and selectively limiting downstream mechanoresponsive signalling, including YAP activation. These effects are consistently observed in both established and patient–derived PDAC models, underscoring their robustness and translational relevance.

Recent studies have highlighted the context–dependent activity of budesonide in PDAC, with pronounced antiproliferative effects in 3D systems but limited activity in conventional 2D cultures^27^. Our findings further suggest that this apparent context dependency may largely reflect differences in the mechanical microenvironment rather than intrinsic drug efficacy, emphasizing the importance of incorporating physiologically relevant stiffness cues into preclinical models. More broadly, our results support the concept that matrix mechanics is a critical determinant of drug response in PDAC^3,4,9,12,24^.

At the nuclear level, our data provide the first evidence, to our knowledge, that budesonide promotes a ‘soft–like’ nuclear state despite persistent mechanical cues, pointing to the nucleus as a key target of its activity. By weakening cytoskeletal–nucleoskeletal coupling, budesonide limits force transmission and attenuates downstream transcriptional responses^41,46,48,51,53^. Indeed, budesonide reduced lamin A/C abundance, induced a higher wrinkling of the nuclear envelope, and constrained nuclear enlargement and deformation induced by stiff substrates. These structural changes were accompanied by cytoplasmic retention of YAP without changes in total protein levels, indicating selective inhibition of mechanoresponsive nuclear signalling rather than generalized protein suppression. Given the emerging role of the nucleus as a mechanosensitive organelle that integrates extracellular forces into transcriptional outputs^32,41–44,51,54^, our findings suggest that budesonide interferes with a mechanotransduction axis linking matrix stiffness to the nucleus and eventually on malignant gene expression programs. Transcriptomic profiling strongly supports this model. Budesonide reversed a broad set of stiffness–induced transcriptional changes associated with proliferation, migration, cytoskeletal remodelling, and aggressive cell states. Thus, the phenotypic effects observed at the cellular and nuclear levels converge with genome–wide transcriptional rewiring, indicating that budesonide does not simply block isolated downstream pathways but resets a broader mechanoadaptive state. This system–level reprogramming may be particularly relevant in PDAC, where stromal stiffening and desmoplasia continuously reinforce tumor cell plasticity and therapeutic resistance^6,7,9–13^.

Our findings further reveal a complex involvement of GR signalling. Glucocorticoid signalling encompasses both canonical GR–dependent transcriptional effects and non–genomic activities^55^, and our data further uncover a previously unappreciated role for GR signalling in regulating PDAC cell mechanics, while also supporting the existence of GR–independent mechanisms underlying budesonide activity. Our results therefore suggest that budesonide engages these pathways in the context of PDAC mechanobiology^28^. Notably, budesonide, similarly to cholesterol, is known to modulate the structural properties of lipid bilayers, including lipid order and packing^56^. By altering plasma membrane fluidity, budesonide may affect membrane organization and receptor clustering, ultimately impairing cell–matrix coupling and mechanosensing^57,58^. While this possibility warrants further investigation, GR silencing experiments uncovered an unappreciated role for GR signalling in regulating PDAC cell behaviour, in particular in cell motility. These results are also supported by the recent evidence that GR regulates ovarian cancer cell motility^59^. Moreover, our results suggest that the effects of budesonide on nuclear organization are largely independent of GR.

Collectively, our data identify matrix stiffness as a dominant regulator of PDAC cell phenotype and establish budesonide as a pharmacological agent capable of functionally uncoupling tumour cells from pathological mechanical cues. Rather than directly targeting oncogenic drivers, budesonide perturbs the biomechanical circuitry that sustains malignant phenotypes, revealing mechanotransduction as a targetable vulnerability in PDAC. Given its established clinical use and favourable safety profile, budesonide represents a compelling candidate for therapeutic repurposing, particularly in combination with treatments targeting genetic or metabolic dependencies.

Several limitations should be acknowledged. Although our platforms capture tuneable stiffness, they do not fully reproduce the multicellular and biochemical complexity of the native PDAC microenvironment. The precise molecular mediators linking budesonide to cytoskeletal and nuclear remodelling remain to be defined, and the relative contribution of GR–dependent versus independent pathways will require deeper genetic and biochemical dissection. Nonetheless, the consistency of our findings across cell lines and patient–derived models underscores the robustness of the observed phenomenon.

In summary, we identify budesonide as a suppressor of stiffness–driven malignancy in PDAC and reveal nuclear mechanotransduction as a targetable vulnerability in this disease. More broadly, our work demonstrates that integrating mechanobiology into cancer pharmacology can uncover actionable therapeutic activities that remain hidden in conventional culture systems.

## METHODS

### Substrate preparation

Polyacrylamide (PAAm) substrates were prepared following established protocols^60^ with modifications to accommodate different surface areas. Glass coverslips, having a diameter of 35 and 50 mm respectively, were silanised using 3-aminopropyltriethoxysilane (APTES, Sigma-Aldrich) for 10 min and glutaraldehyde 0.5% for 10 min. Every step accounted for accurate washing with double-distilled water to remove excess reagent.

PAAm hydrogels were prepared by mixing acrylamide and N,N′-methylenebisacrylamide (Sigma-Aldrich) in PBS 1×. The final concentrations used were 2.5% acrylamide/0.15% bis-acrylamide, 6% acrylamide/0.15% bis-acrylamide and 10% acrylamide/0.13% bis-acrylamide. The crosslinking was triggered using ammonium persulfate (APS, Sigma-Aldrich) 10% at 1/100 final concentration and N,N,N′,N′-tetramethylethylenediamide (TEMED, Sigma-Aldrich) at 1/1000 final concentration. The prepared gels were pre–tested with a stress–driven rheometer (MCR302, Anton Paar), using a parallel plate geometry with an upper plate diameter of 25 mm. Dynamic strain tests at constant temperature (37 °C) were performed both at a constant frequency (1 Hz), applying a strain ranging from 1 to 100%, and through a frequency sweep from 0.1 Hz to 10 Hz, resulting in quantification of the complex shear modulus G*. Young’s modulus was then calculated as indicated in **Equation 1**.

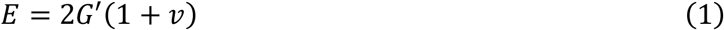

where *v* is the Poisson’s number, assumed to be equal to 0.5 for PAAm^61^. Due to the elastic behaviour of the material and due to the negligible contributions of the loss moduli, only the storage modulus G’ was considered in the equation. Rheometer testing yielded a final stiffness of the hydrogels of 0.2, 10 and 27 kPa in Young Modulus respectively. Measurements to assess the consistency of the obtained formulations in terms of Young’s modulus were performed at each gel preparation (Supplementary Table 3 and Supplementary Fig.1A). To cast the PAAm substrates, the activated solution was applied to the treated side of the coverslips mounted to the appropriate dishes and covered with a circular coverslip (10, 20, or 50 mm in diameter) before allowing polymerization at 37 °C for 20 min. After polymerization, the top coverslips were removed, resulting in cylindrical hydrogels with an approximate thickness of 220 µm. The substrates were then soaked with a solution of 50% Penicillin/Streptomycin (Microgem) and 10% Amphotericin-B (Sigma-Aldrich) in sterile PBS 1× for a minimum of 24 h at 4 °C, followed by exposure to UV germicidal lamp (265 nm) for at least 1 h two times for complete sterilization. To allow cell culture, the substrates were functionalized with type I collagen. Specifically, a 0.2 mg/mL solution of bifunctional crosslinker sulfosuccinimidyl 6-(4′-azido-2′-nitrophenylamino) hexanoate (sulfo-SANPAH, ThermoFisher) in sterile–filtered double–distilled water was applied to the gel surface and photoactivated under 365 nm UV light for 10 min. After being washed with sterile PBS, the substrates were coated with a 50 µg/mL collagen solution (Bovine Type I Collagen solution, Sigma-Aldrich) in sterile–filtered double–distilled water at 37 °C for 1 h. The substrates were then washed with sterile PBS and used for cell culture within 2 days.

### Cell lines, culture conditions, and treatments

The human pancreatic cancer cell line PANC-1 (ATCC, CRL-1469™) and two patient–derived PDAC lines (PDAC#253 and PDAC#354), previously isolated and characterized from independent pancreatic ductal adenocarcinoma samples^39,40^, were used in this study. All cultures were routinely screened to confirm the absence of Mycoplasma contamination and were maintained below passage 28.

PANC-1 cells were grown in DMEM (Invitrogen) supplemented with 10% fetal bovine serum (Invitrogen), L-Glutamine (Invitrogen), and Penicillin/Streptomycin (Invitrogen). PDAC#253 and #354 were maintained in a customized PDAC growth medium consisting of RPMI 1640 with GlutaMAX™ (Invitrogen), 10% South American FBS (Euroclone), Penicillin/Streptomycin (Invitrogen), and Normocin (InvivoGen).

For treatment experiments, PANC-1, PDAC#253 and PDAC#354 cells were initially seeded at 15 × 10³ cells/cm² on gelatin–coated dishes, at day 0 and day –1, respectively, and treated with budesonide (20 µM), dexamethasone (20 µM), or no treatment. After 3 days, cells were trypsinised and reseeded at 2.5 × 10³ cells/cm² on bioengineered substrates, with or without drugs for additional two days. Following treatments, cells were either fixed with 4% paraformaldehyde (PFA) or processed for downstream applications.

Budesonide (Sigma-Aldrich) was dissolved in DMSO, while dexamethasone (Sigma-Aldrich) was dissolved in water at 10 mM.

### Generation of *NR3C1* KD PANC-1 cells

Stable knockdown of NR3C1 in PANC-1 was achieved by transduction with lentiviral vectors encoding shRNAs targeting NR3C1 along with a puromycin resistance cassette^38^. Cells transduced with an empty vector (shEmpty) served as negative control. After 48 h, puromycin selection (5 µg/ml) was applied for 7 days to obtain stable populations.

### Cell proliferation assays

To evaluate proliferation, PANC-1 cells were treated as described above and then labelled using the Click-iT EdU Flow Cytometry Assay Kit (Invitrogen), according to the manufacturer’s instructions. After fixation and staining, samples were analysed by flow cytometry (FACS-ARIAIII, Becton-Dickinson). The growth rate of the cells was extracted from time–lapse microscopy experiments (LEICA STED-SP5). Cell number for each video was assessed at the starting frame (nC_T0_) and at 48 h (nC_T48_), determining the growth rate through **Equation 2** which describes the number of replications per day.

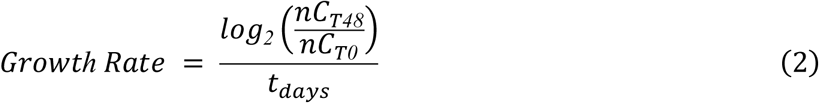

### Immunostaining

Upon treatments, PANC1, PDAC #253 and PDAC #354 cells were fixed with PFA 4% for 10 min. For analysis of cytoskeleton markers, cells were then permeabilized with TRITON 0.1% for 10 min at room temperature (RT) and then incubated with a blocking solution of BSA 5%/TRITON 0.1% for 1 h. Primary antibodies were incubated overnight at 4 °C, followed by the respective secondary antibodies (Alexa Fluor Molecular Probes). Images were acquired with a Nikon A1 confocal microscope. The NIS Element C (Nikon, Tokyo) software was used for image acquisition/elaboration. For focal adhesions’ area quantification, paxillin was stained with rabbit anti–paxillin monoclonal antibody followed by incubation with an appropriate secondary antibody. Cytoskeletal actin fibres were stained with Alexa Fluor 647-conjugated phalloidin. Images were acquired using a confocal laser scanning microscope (LSM-900, Carl Zeiss) equipped with two diode laser lines (green channel, λ_excitation_ = 488 nm, λ_emission_ = 500 – 700 nm, for paxillin, and deep red channel, λ_excitation_ = 633 nm, λ_emission_ = 651 – 720 nm for phalloidin). The images were collected as single snaps using a 40× C-Apochromat korr water immersion objective M27 (NA = 1.2) at an image resolution of 2048 × 2048 pixels with a 2× zoom factor.

For YAP translocation investigations, cell membranes were stained with rhodamine–wheat germ agglutinin (WGA) before permeabilization. Then, permeabilization was achieved using a solution of TRITON 0.3% in PBS 1× buffer for 15 min, followed by a blocking solution of BSA 3%/TRITON 0.1% in PBS 1× buffer for 1.5 h. Cells were then incubated with rabbit anti–YAP polyclonal antibody followed by incubation with an appropriate secondary antibody. Images were acquired using a confocal laser scanning microscope (LSM-710, Carl Zeiss) equipped with three diode laser lines (green channel, λ_excitation_ = 488 nm, λ_emission_ = 495 – 544 nm for the nuclei, red channel, λ_excitation_ = 555 nm, λ_emission_ = 546 – 594 nm for cellular membrane, and deep red channel λ_excitation_ = 647 nm, λ_emission_ = 651 – 720 nm for YAP). The images were collected at 1 µm Z-spacing using a 40× C-Apochromat korr water immersion objective M27 (NA = 1.2) at an image resolution of 1024 × 1024 pixels with a 2× zoom factor.

For investigation of the nuclear laminae, cells were permeabilized with a solution of Glycine 100 mM/TRITON 0.3% in PBS 1× buffer for 15 min, then a blocking solution of BSA 3%/TRITON 0.1% in PBS 1× buffer was applied for 1.5 h. Cells were then incubated with rabbit anti–lamin B1 polyclonal antibody and subsequently with mouse anti–lamin A/C monoclonal antibody, followed by incubation with the respective secondary antibodies. Images were acquired using a confocal laser scanning microscope (LSM-900, Carl Zeiss) equipped with three diode laser lines (blue channel, λ_excitation_ = 405 nm λ_emission_ = 410 – 475 nm for the nuclei, green channel, λ_excitation_ = 488 nm, λ_emission_ = 500 – 700 nm, for lamin A/C, and red channel, λ_excitation_ = 546 nm, λ_emission_ = 410 – 617 nm for lamin B1). The images were collected at 1 µm Z-spacing using a 40× C-Apochromat korr water immersion objective M27 (NA = 1.2) at an image resolution of 1024 × 1024 pixels with a 2× zoom factor. The list of primary and secondary antibodies with the respective dilution and incubation time is shown in Supplementary Table 4.

### Immunofluorescence analysis

For spreading area quantification, the cell area on the adhesion plane slice was directly measured using FIJI ImageJ (NIH software). Focal adhesions area was calculated by isolating peripheral focal adhesion from paxillin staining. Filtering of the image, based on a custom written ImageJ routine, was performed prior to manual selection of isolated peripheral adhesions overlapped with phalloidin staining and identified as belonging to ventral stress fibres, which are inherently related to nucleoskeletal mechanotransduction through actin cap connection.

YAP localization was analysed using IMARIS (IMARIS, Oxford Instruments) for 3D rendering and quantification. 3D models of individual cells were generated using the "Surface" function on WGA and SYTOX fluorescence signals obtained from confocal imaging, enabling the isolation of a nuclear and a cellular compartment. Cytoplasmic volume was obtained from **Equation 3**. YAP intensity was evaluated within the produced compartments and extracted as mean grey values (MGVs), which represent the average concentration of YAP at each reconstructed voxel. MGV for cytoplasm was extracted from **Equation 4**. This representation was chosen to avoid corruption of the dataset due to the dimensional heterogeneity of PANC-1 cells within the same condition.

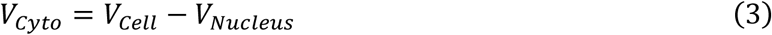

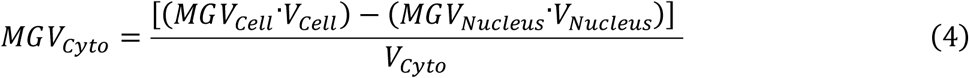

where V_Cyto_, V_Cell_ and V_Nucleus_ are the volumes of the cytoplasm, cell and nucleus respectively; and MGV_Cyto_, MGV_Cell_ and MGV_Nucleus_ are the mean grey values of YAP signal within cytoplasm, cell and nucleus compartments. Localization of YAP was quantified by means of N/C intensity ratio, as described in **Equation 5**.

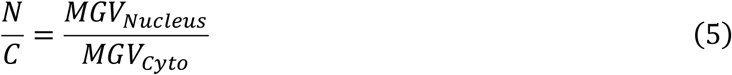

To extract data for lamin A/C and lamin B1 expression, the slice containing maximal area extension for the nuclei of every imaged cell was singled out from the z-stack. Then, ROIs were determined by constructing a selection mask through Threshold function on the lamin B1 signal. Within the selected ROIs, MGVs for the lamin A/C and lamin B1 signals were extracted. Nuclear wrinkling for lamin B1 was evaluated by adapting an existing protocol to the acquired images^41^. Sum projections of complete nuclear z-stacks for each condition were provided to a customized ImageJ routine extracted from literature^38^. A selection mask allowed for recording of mean intensity of sum images and their aspect ratio. Contour tracking for the nuclei was executed using FeatureJ via a Sobel–based edge detection. Quantification of wrinkling was determined through a wrinkling index, defined as the percentual area coverage on the nucleus detected by the Sobel-based edge detection, visualized as the white pixels of the binarized and FeatureJ–processed image.

### Cell motility

Cell motility experiments were performed by time–lapse microscopy (LEICA, STED-SP5). Transmission images were collected at 10-11 min intervals for at least 48 h, with 254-288 frames analysed per cell. Cell positions in each frame were manually tracked using ImageJ, via the Manual Tracking plugin. After the complete positions set was obtained, the data were converted in micron and cell velocity and mean–square displacement (MSD) were calculated as described in **Equation 6-7** through a routine derived from literature^62^.

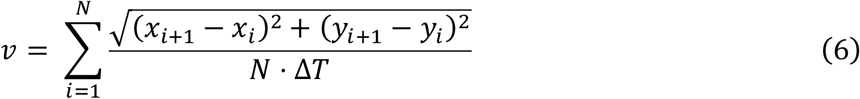

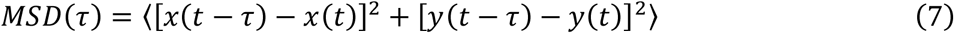

Where x_i_ and y_i_ are the spatial coordinates in 2D space for each cell at the i-th frame, ΔT is the time in minutes occurring between two subsequent frames, N is the total number of acquired frames, t is the time variable, τ is the lag time and 〈 〉 indicates the time average operator. In principle, only isolated cells exhibiting single–cell migration without division or cell–cell contact were included. Due to the prominent replication capabilities of PANC-1, PDAC #253 and PDAC #354 cells, the selection criterion was adjusted to amplify the dataset without disrupting the validity of the executed tracking. Specifically, in cases where only a single division occurred in the complete timelapse acquired, tracking was continued following the first daughter cell that remained within the field of view and displayed stable substrate adhesion. This allowed for the inclusion of a larger number of cells and an extended dataset.

### Western blot

Cells were lysed in a buffer containing 20 mM Tris (pH 8), 150 mM NaCl, 10 mM EDTA, 1% Triton X-100, 10% glycerol, and 1 mM zinc acetate. Equal amounts of protein were separated by SDS-PAGE and transferred to PVDF membranes using the Trans-Blot SD Semi-Dry Transfer Cell (BIO-RAD). Membranes were incubated overnight at 4 °C with primary antibodies (Supplementary Table 4), followed by HRP-conjugated secondary antibodies. Protein detection was performed with ECL reagents (EuroClone), and band intensities were quantified using ImageLab software.

### AFM

The mechanical properties of PANC-1 cells were assessed using a JPK NanoWizard II AFM, combined with an optical microscope for precise positioning of tips and samples. Soft cantilevers (SAA-SPH-5UM, spring constant 0.171 N/m) were employed to probe cell mechanics in force spectroscopy mode. For each measurement, 64 force curves were collected over a 2 × 2 μm² surface area. Young’s modulus was derived from each force curve within the force map using JPKSPM data processing software. The Hertzian model was applied to calculate Young’s modulus for each curve, generating 64 values per map, according to the relation in **Equation 8**, which describes the relationship between the indentation δ and the applied force F for an infinitely hard spherical tip of radius r contacting a soft planar surface.

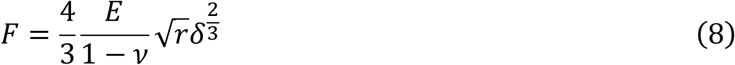

where E is Young’s modulus and ν is the Poisson ratio, assumed equal to 0.5. Results showed the average value calculated from each force curve as a representative modulus of each measured force.

### RNA extraction and Sequencing

For RNA–seq experiments, PANC-1 cells were cultured on gelatin–coated plates in the presence or absence of budesonide (20 µM) for three days, then reseeded at 2.5 × 10³ cells/cm² on engineered platforms and maintained for two additional days with medium refreshed ± drug. Total RNA was isolated using the RNeasy kit (Zymo Research). RNA sequencing was carried out by Genomix4Life (https://www.genomix4life.com/en/bioinformatic_technologies.html) using the Illumina NovaSeq 6000 platform. Raw sequencing reads were aligned to the GenCode reference genome (Release 48, GRCh38.p14). Gene expression levels were quantified with FeatureCounts (version 2.0.3). A count matrix including all expressed genes across all samples was generated in R, and normalization together with differential expression analysis was performed using the Bioconductor package DESeq2 (significance threshold p < 0.05). Normalization was achieved by dividing counts by sample–specific size factors, calculated as the median ratio of each gene’s read count relative to its geometric mean. Gene Ontology (GO) enrichment was performed with the DAVID tool (https://david.ncifcrf.gov/#). Furthermore, pathway analysis was carried out in R for KEGG, Biological Process (BP), and Reactome pathways. KEGG enrichment was conducted using the clusterProfiler package in R, ranking genes by differential expression values. For Reactome analysis, the reactome.db package was applied to map differentially expressed genes to relevant pathways. Only genes with an adjusted p-value ≤ 0.05 were considered for pathway enrichment. Results from differential expression and pathway analyses (KEGG, BP, and Reactome) were integrated to obtain a broader understanding of the biological processes involved.

### Statistical analysis

The data were represented as mean ± standard deviation (SD) or ± standard error of the mean (SEM), as well as through boxplots or dotplots. The number of independent experiments is shown as “N=” and the total sample size is provided in each figure legend (n). Normality of the datasets acquired was determined using the Shapiro-Wilk test (<5000 datapoints) while the statistical comparisons were either performed with the two-tailed Student’s paired t-test for normal distribution pairs or the Mann Whitney U-test in all other cases. Choice of pooled variance t-test or Welch correction t-test was determined by execution of an F-test on the analysed sample pair sets. P-values <0.05 denote statistically significant differences. Data derived from statistical analysis are explicitly included in Supplementary Tables 5-15. Data representations, including graphs and boxplots, were generated using either Microsoft Excel, OriginPro or RStudio software, specifically version 1.1.463 from RStudio, Inc., accessible at https://www.rstudio.com/.

## DECLARATIONS

### Author contributions

V.P. and C.D. conceived and designed the study. V.P. and C.D. supervised the study. S.M., M.C. and E.I. performed experiments and analysed data. E.I. performed RNA sequencing analysis. A.A. prepared PAAm substrates and analysed migration data. C.F. analysed focal adhesions data. B.S. JR and E.L., provide reagents and gave conceptual advice. L.A. and G.C., discussed and provide conceptual advice. E.J.P., G.M., P.A.N. contributed to data interpretation and scientific discussion. S.M., M.C., E.J.P., G.M., V.P. and C.D. wrote the manuscript with input from all authors. All authors read and approved the manuscript.

### Availability of data

RNA-Seq data have been deposited to GEO repository (Submission ID: GSE330775). Raw data and analysed data can be supplied by authors upon reasonable request.

## Acknowledgments

We thank Salvatore Arbucci, Vincenzo Mercadante and members of the Integrated Microscopy and FACS Facilities of the IGB-ABT, CNR. We acknowledge Prof. Giannino Del Sal for kindly providing the NR3C1shRNA construct. We also acknowledge the PhD school in Precision Medicine of the University of Palermo, Italy.

---

**Supplementary Figure 1.**
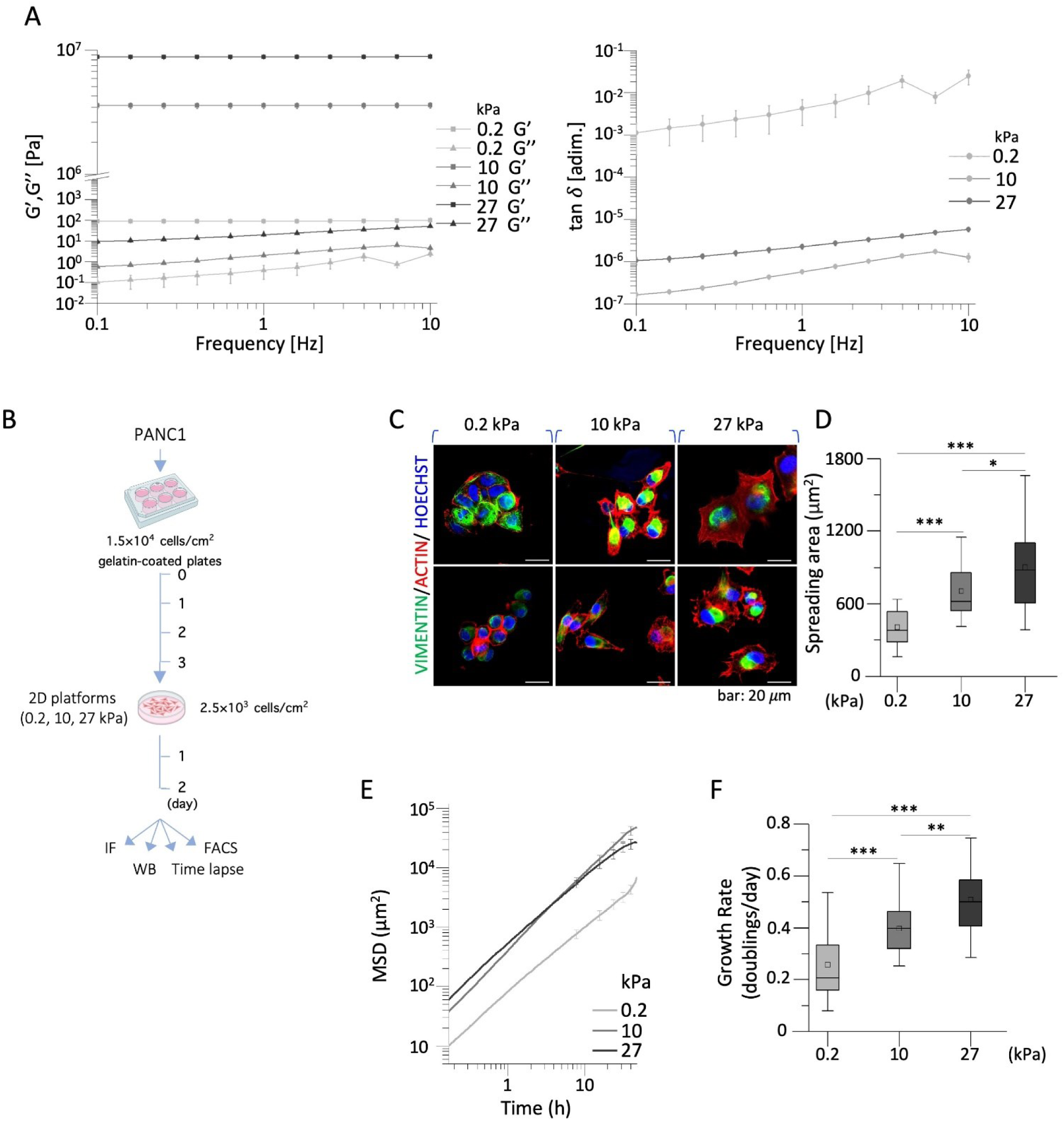
PANC-1 cells adapt their mechanical and biological properties to increasing substrate stiffness. **A** Rheological properties of PAAm substrates. Storage modulus (G′) and loss modulus (G″) are plotted as a function of frequency and measured at 37 °C (*left*). Corresponding damping factor (tan δ = G″/G′) is plotted over the same range (*right*), indicating the dominance of the elastic response in PAAm behaviour. **B** Schematic representation of the experimental procedure. PANC-1 cells were plated at day 0 on gelatin-coated plates at 1.5×10^4^ cells/cm^2^. After three days in culture, cells were detached with trypsin and plated at 2.5×10^3^ cells/cm^2^ on the bioengineered platforms at different stiffness (0.2, 10, and 27 kPa) for two days. **C-D** Representative confocal images of vimentin (green) and F-actin filaments (red) staining in PANC-1 cells (C), and spreading area analyses (D, n=40). Nuclei were stained with HOECHST (blue). **E** MSDs of PANC-1 cells cultured on 0.2, 10, and 27 kPa substrates and acquired through a 48 hrs timelapse. **F** Quantification of cell growth rate on 0.2, 10, and 27 kPa substrates from time-lapse microscopy analyses (n=20). Quantitative data are presented as box plots (N=3). Significant statistical differences are reported: * p < 0.05; ** p < 0.01; *** p < 0.001. The complete statistical analysis executed on shown datasets can be found in Supplementary Table 6.

**Supplementary Figure 2.**
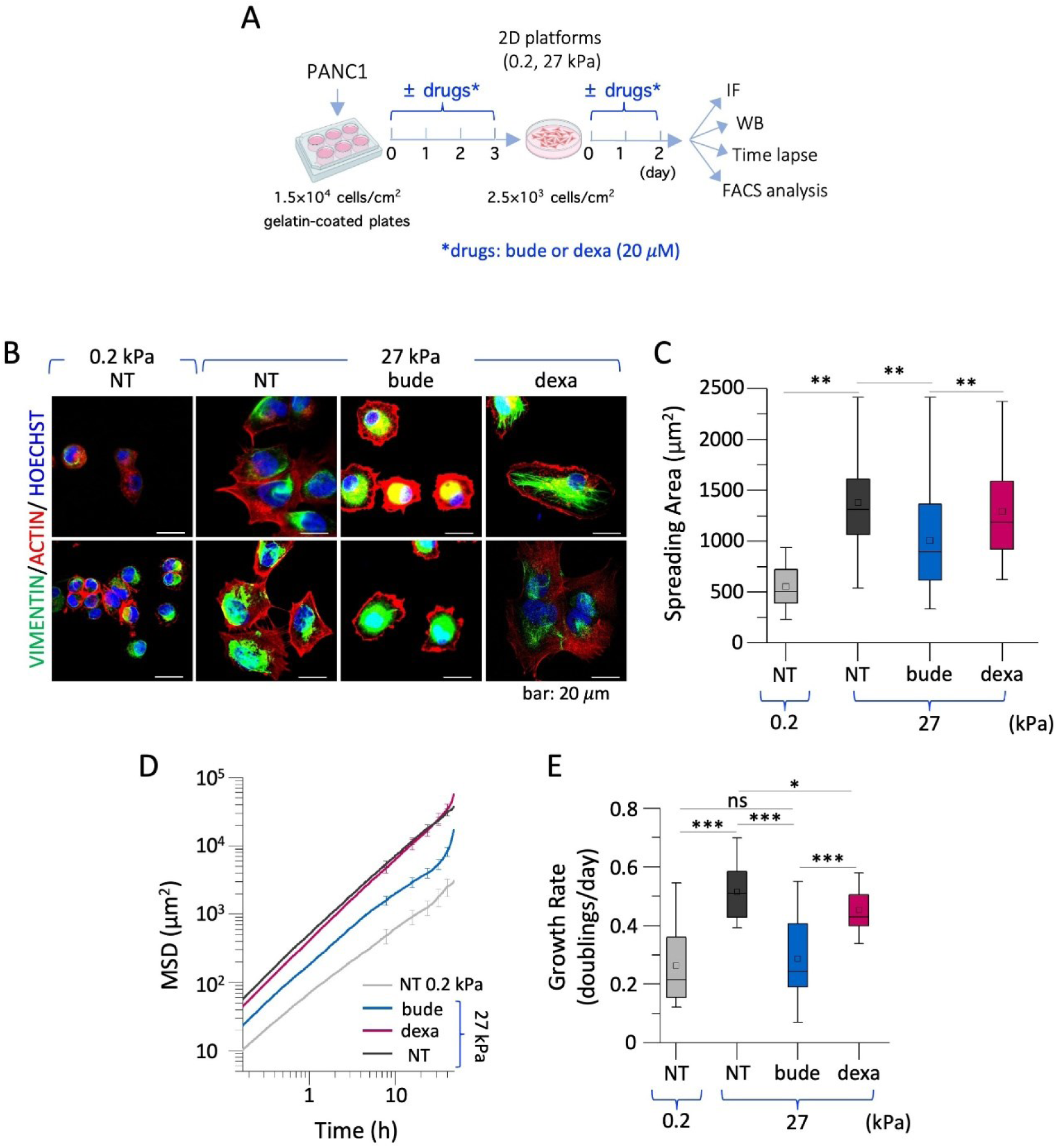
Budesonide effects on PANC-1 cell behaviour under stiff conditions**. A** Schematic representation of the experimental procedure. PANC-1 cells were plated at day 0 on gelatin-coated plates at 1.5×10^4^ cells/cm^2^, ± budesonide (20 µM) or dexamethasone (20 µM). After three days in culture, cells were detached with trypsin and plated at 2.5×10^3^ cells/cm^2^ on the stiffer 27 kPa substrate for two days, ± the same treatments. Control NT cells were also plated on the 0.2 kPa substrate. **B-C** Representative confocal images of vimentin (green) and F-actin filaments (red) staining in treated PANC-1 cells (B), and spreading area analyses (C, n=40). Nuclei were stained with HOECHST (blue). **D** MSDs of PANC-1 cells under the indicated treatments and stiffness conditions. **E** Quantification of growth rate of treated PANC-1 cells extracted from time-lapse microscopy (n=20). Quantitative data are presented as box plots (N=2-3). Significant statistical differences are reported: * p < 0.05; ** p < 0.01; *** p < 0.001. The complete statistical analysis executed on shown datasets can be found in Supplementary Table 8.

**Supplementary Figure 3.**
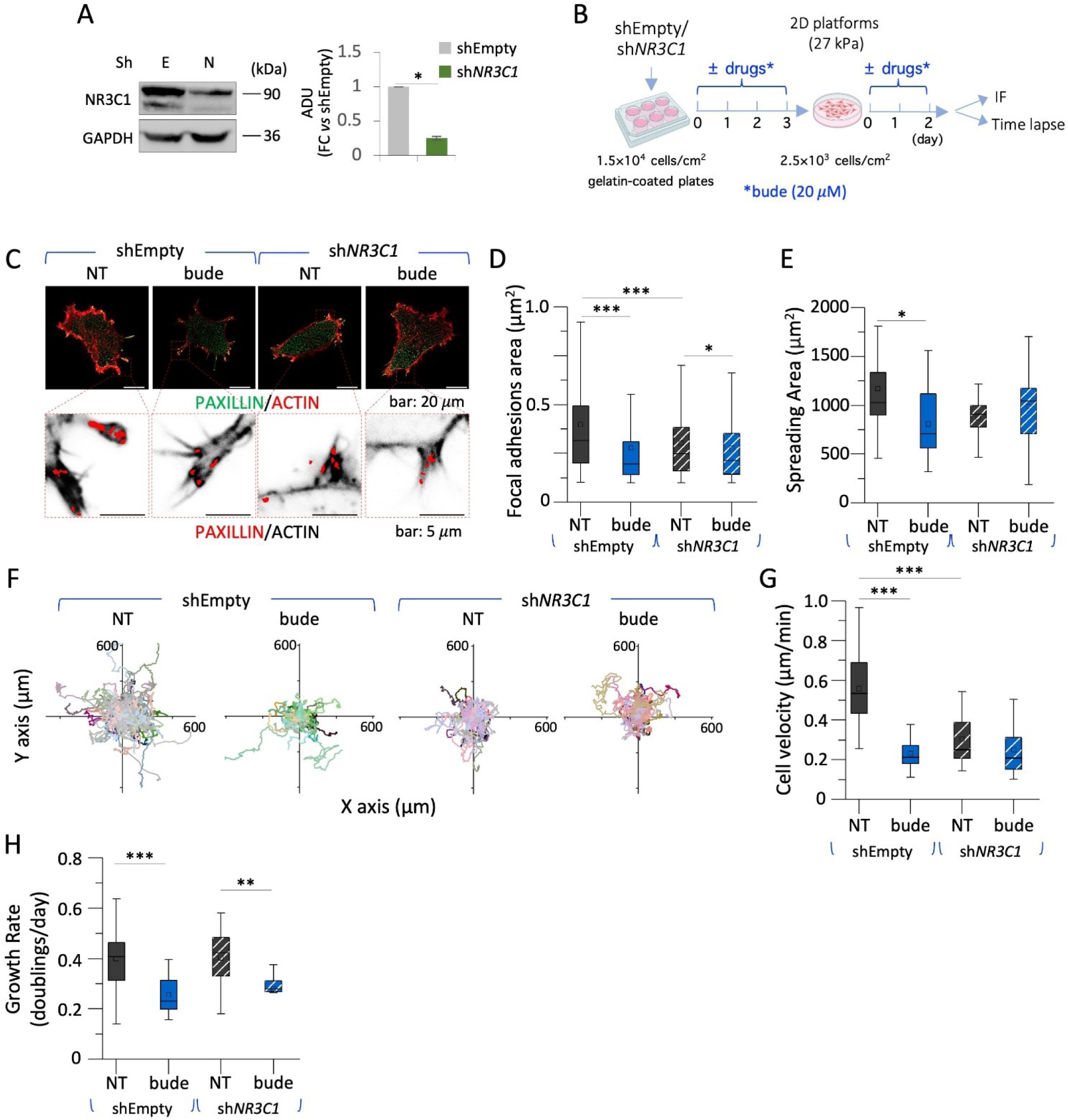
Study of the GR involvement into the mechanism of action of budesonide. **A** Representative western blot analysis (*left*) and densitometric quantification (ADU; *right*) of GR level of expression in *^shNR3C1^*PANC-1 cells versus *^shEmpty^*PANC-1 cells. **B** Schematic representation of the experimental procedure. PANC-1 silenced cells (sh*NR3C1*/shEmpty) were plated at day 0 on gelatin-coated plates at 1.5×10^4^ cells/cm^2^, ± budesonide (20 µM). After three days in culture, cells were detached with trypsin and plated at 2.5×10^3^ cells/cm^2^ on the stiffer 27 kPa substrate for two days, ± the same treatment. **C-E** Representative confocal images of *^shEmpty^*PANC-1 and *^shNR3C1^*PANC-1 cells, showing paxillin (green) and F-actin filaments (red) staining with focal adhesions highlighted in yellow (*upper*-C). The corresponding zoomed inverted contrast views with focal adhesions’ ROIs selected from paxillin (red) and F-actin (black) are shown underneath (*lower-*C). Focal adhesion area (D) and spreading area analyses (E) are shown (n=30). **F-G** Plotted single-cell trajectories of *^shEmpty^*PANC-1 and *^shNR3C1^*PANC-1 cells upon treatment with budesonide (F) and single-cell velocity analyses in time lapse microscopy (G, n=100) with **H** Quantification of cell growth rate of *^shEmpty^*PANC-1 and ^s*hNR3C1*^PANC-1 cells upon treatment with budesonide (n=20). Quantitative data are presented as box plots (N=3). Significant statistical differences are reported: * p < 0.05; ** p < 0.01; *** p < 0.001. The complete statistical analysis executed on shown datasets can be found in Supplementary Table 9.

**Supplementary Figure 4.**
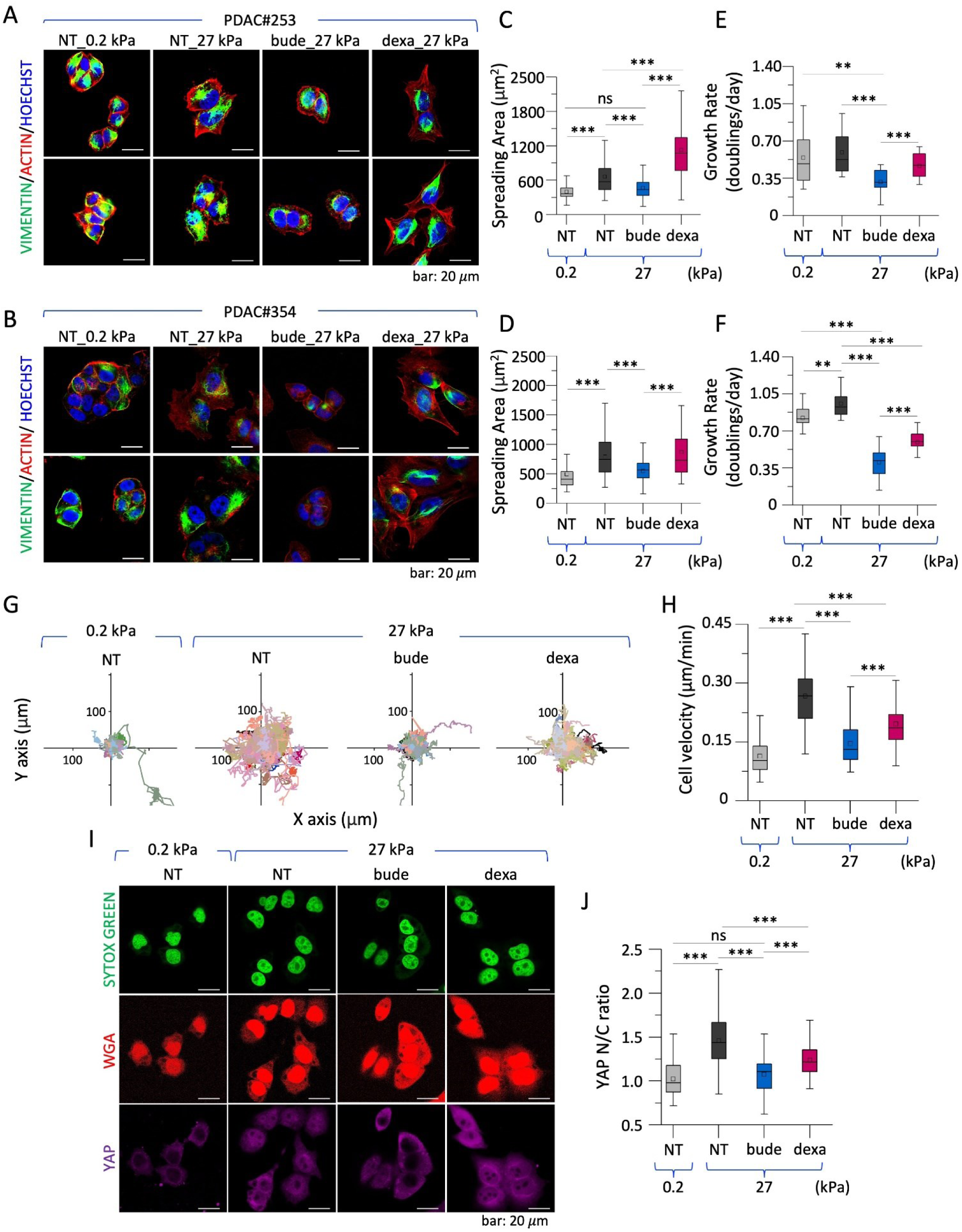
Budesonide reduces cytoskeletal organization, proliferation, and migration in patient-derived PDAC cells. **A–D** Representative confocal images of vimentin (green) and F-actin filaments (red) staining in PDAC#253 (A) and #354 cells (B), upon culture on the bioengineered substrates (0.2 and 27 kPa) and treated ± budesonide or dexamethasone (20 µM). Nuclei were stained with HOECHST (blue). The images are shown alongside the corresponding spreading area analyses for PDAC#253 (C) and PDAC#354 (D, n=110/60). **E–F** Quantification of cell growth rate from time-lapse microscopy analyses of PDAC#253 (E) and PDAC#354 (F) upon treatment (n=20). **G-H** Plotted single-cell trajectories of PDAC#354 cells (G) and single-cell velocity analyses in time lapse microscopy (H, n=100) treated as described above. **I-J** Representative confocal images (I) of YAP (purple) and WGA-Rhodamine (red) staining (*left*), with the corresponding YAP N/C ratio intensity analyses (J, n=60), in PDAC#354 cells treated as described above. Nuclei were stained with SYTOX green (*green*). Scale bar: 20 µm. Quantitative data are presented as box plots (N=2). Significant statistical differences are reported: * p < 0.05; ** p < 0.01; *** p < 0.001. The complete statistical analysis executed on shown datasets can be found in Supplementary Table 11 and 12.

**Supplementary Figure 5.**
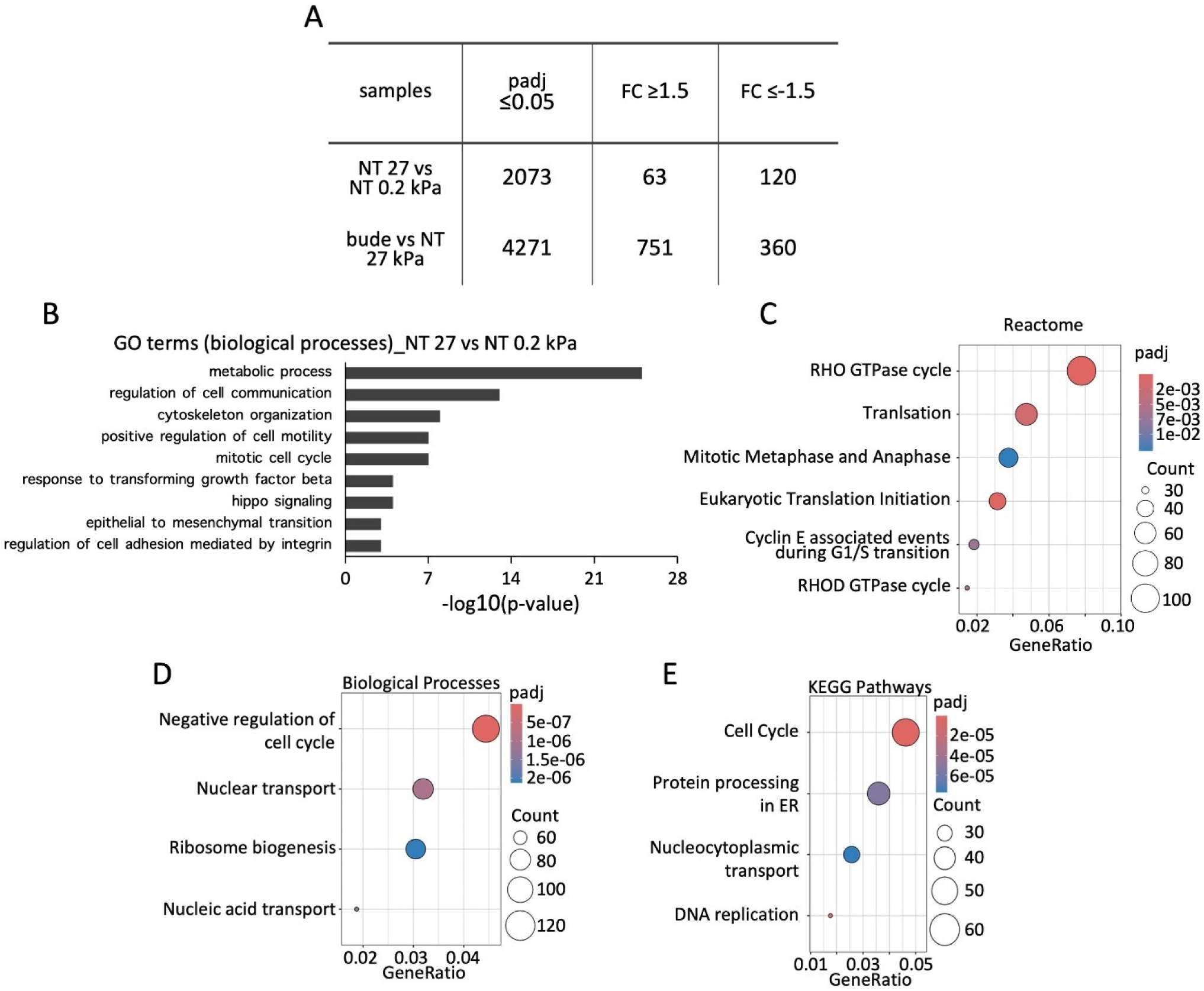
Transcriptomic analysis of stiffness- and budesonide-regulated genes in PANC-1 cells. **A** Table showing the number of deregulated genes in the comparison between NT PANC-1 cells plated on 27 *versus* 0.2 kPa substrates; and between budesonide-treated *versus* NT PANC-1 cells plated on 27 kPa substrate (padj< 0.05; FC <-1.5 and FC > 1.5). **B** Gene Ontology GO analysis (Biological Process) of deregulated genes (padj < 0.05) between NT PANC-1 cells plated on 27 *versus* 0.2 kPa substrates **B-C** Enrichment data analysis of differentially expressed genes for Reactome, in the comparison between NT cells plated at 27 *versus* 0.2 kPa substrates. **D-E** Enrichment data analyses of differentially expressed genes for biological processes (**D**) and KEGG pathways **(E**) specifically modulated by budesonide in PANC-1 cells seeded on 27 kPa substrate.

**Supplementary Figure 6.**
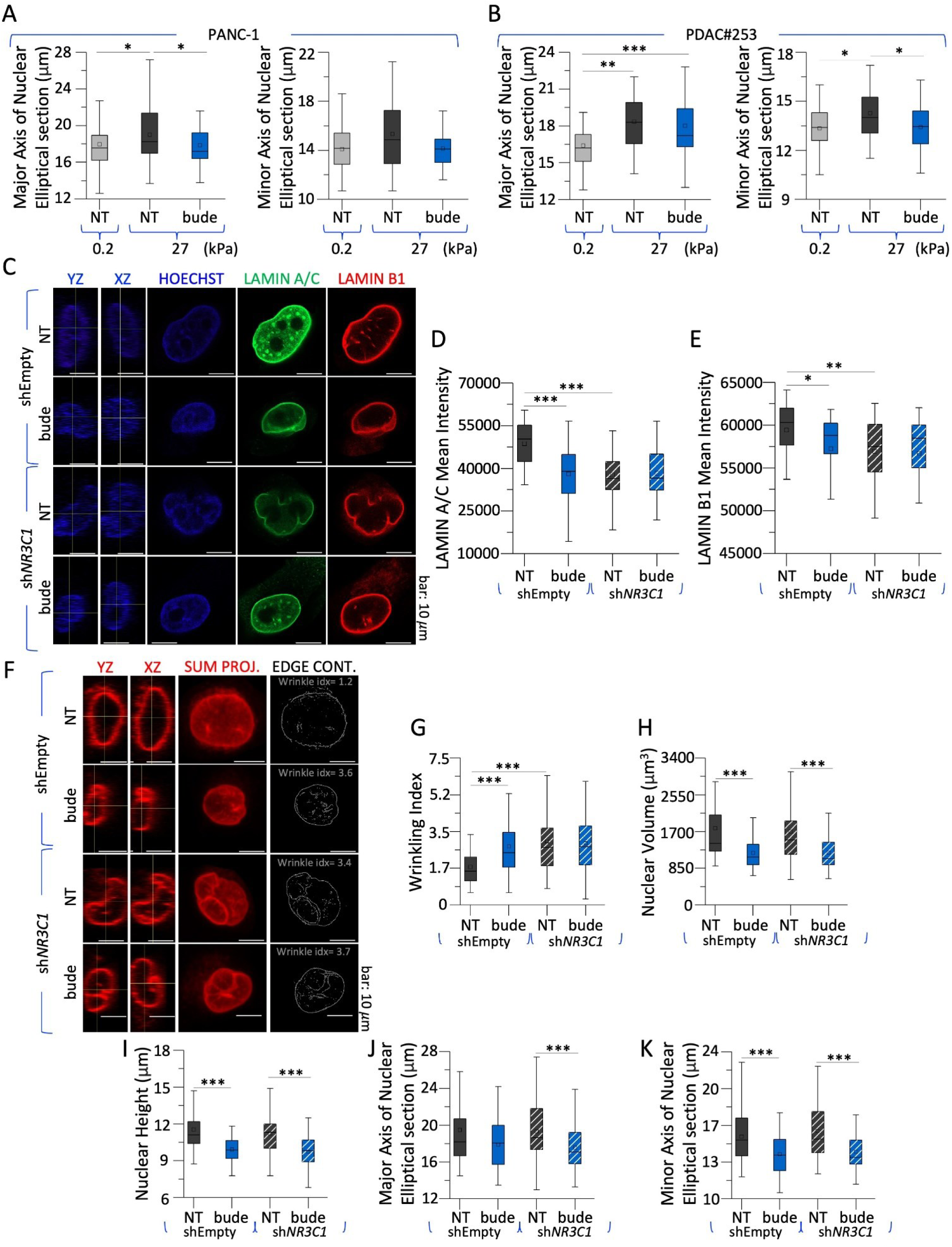
Budesonide impacts nuclear size and shape through GR-independent mechanisms in PANC-1 cells**. A-B** Quantitative morphometry of PANC-1 (A) and PDAC#253 (B) cells nuclei, obtained by measurement of the bounding box encompassing the nuclear signal extracted from HOECHST staining, shown as box plots. **C-E** Representative confocal images (A) of lamin A/C (green) and lamin B1 (red) in *^shEmpty^*PANC-1 and *^shNR3C1^*PANC-1 cells plated on the 27 kPa substrate and treated ± budesonide (20 µM). Nuclei were counterstained with HOECHST (blue). The orthogonal projections in the YZ and XZ planes of representative nuclei (*left*) are shown (C) alongside with the quantifications of lamin A/C (D) and lamin B1 (E) mean fluorescence intensities. **F-H** Representative confocal images of the sum projection, alongside the orthogonal projections in the YZ and XZ planes for lamin B1 (F; red). The representative nuclei are accompanied by edge contour representations extracted using Sobel-based edge filtering, with the wrinkling index quantification (G) and the corresponding nuclear volume (H). **I-K** Quantitative morphometry of *^shEmpty^*PANC-1 and *^shNR3C1^*PANC-1 cells nuclei, obtained by measurement of the bounding box encompassing the nuclear signal extracted from HOECHST staining, shown as box plots. Scale bar: 10 µm. Quantitative data are presented as box plots (n=35, N=3). Significant statistical differences are reported: * p < 0.05; ** p < 0.01; *** p < 0.001. The complete statistical analysis executed on shown datasets can be found in Supplementary Table 15

**Supplementary Table 3.** Rheological measurements obtained from an amplitude sweep test at fixed frequency (1 Hz). The measurements were repeated at each preparation for every polyacrylamide solution formulation and the obtained values allowed for extraction of the derived Young Modulus. The damping factor <<<1 also allows for the fulfilment of the elastic or near-elastic hypothesis of the material used to produce the substrates used for mechanical conditioning.

| **Amplitude Sweep at 1 Hz** |  |  |  |  |
| --- | --- | --- | --- | --- |
| **Formulation** | **Storage Modulus G'**<br>[Pa] | **Loss Modulus G''**<br>[Pa] | **Damping Factor tanδ**<br>[adim.] | **Calculated Young Modulus E**<br>[kPa] |
| 2.5% acrylamide / 0.15% bis-acrylamide | 71.12 ± 10.62 | 0.60 ± 0.41 | 0.01 ± 4.22E-03 | 0.21 ± 0.03 |
| 6% acrylamide/0.15% bis-acrylamide | 3457.16 ± 164.12 | 2.13 ± 0.34 | 6.15E-04 ± 7.54E-05 | 10.37 ± 0.50 |
| 10% acrylamide/0.13% bis-acrylamide | 9044.77 ± 241.26 | 16.46 ± 1.67 | 1.82E-03 ± 1.50E-04 | 27.13 ± 0.72 |

**Supplementary Table 4.**
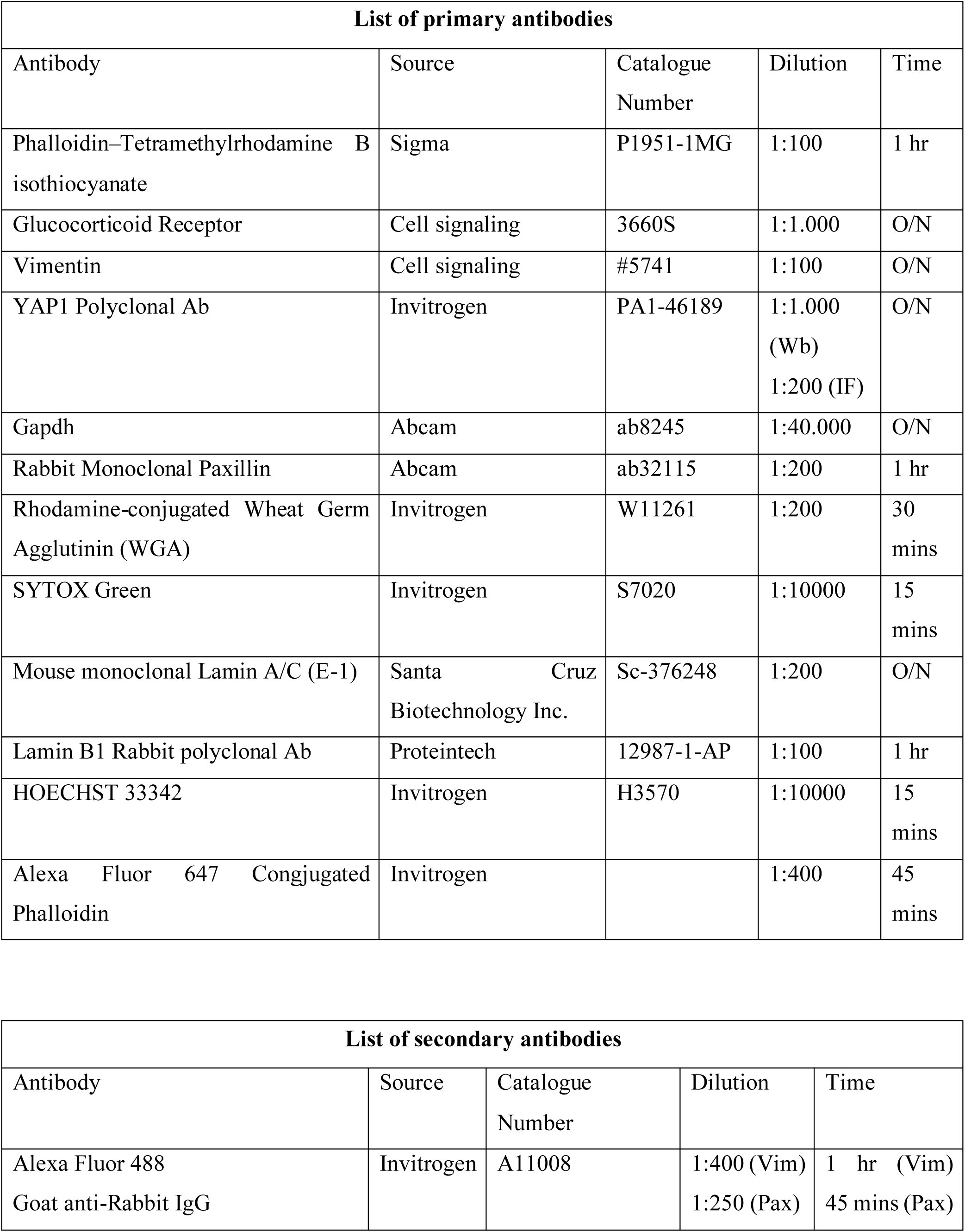
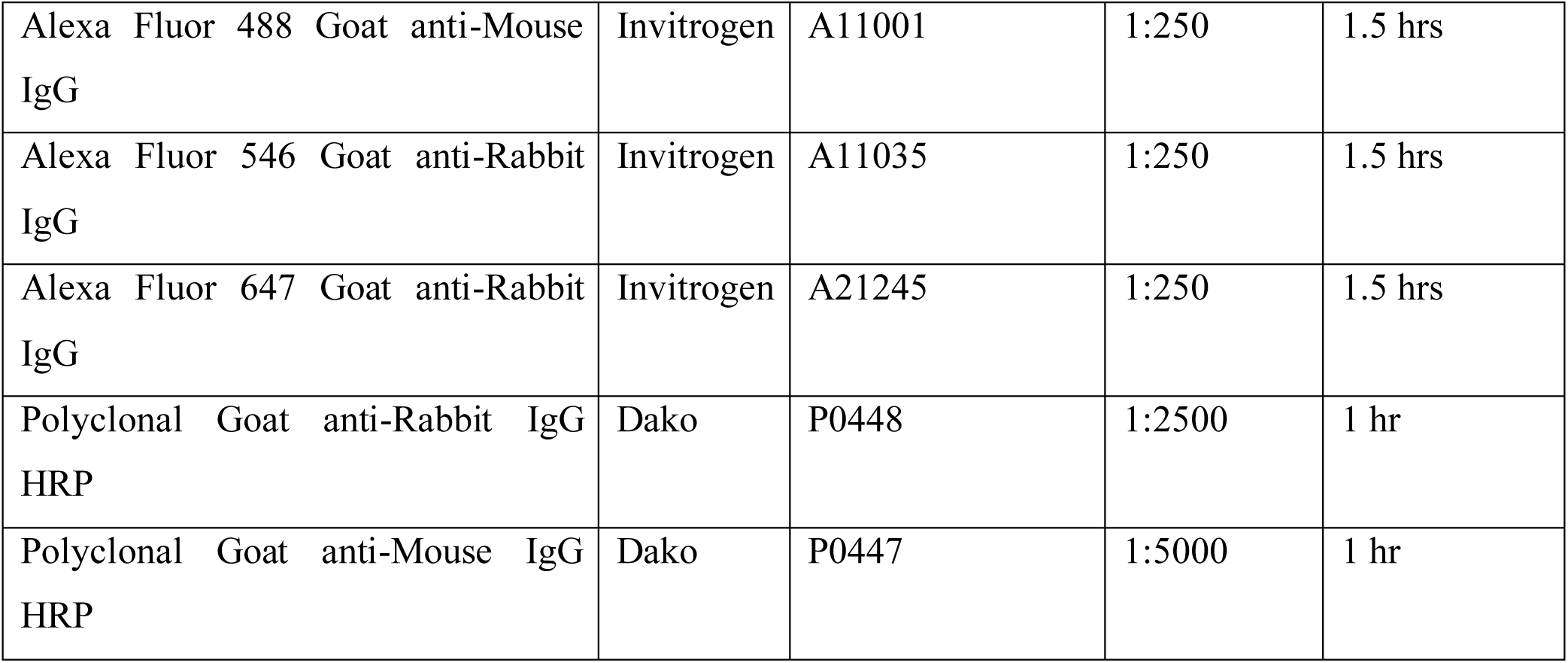
List of the primary antibodies used in IF staining and WBs, as well as their associated secondary antibodies. The table indicates the source producer of the antibodies and their corresponding catalogue number, as well as the protocol used to stain the samples analysed.

**Supplementary Table 5.** Complete statistical analysis for PANC-1-related data shown in Figure 1. For each condition in all the experimental datasets produced (0.2 kPa, 10 kPa and 27 kPa) the normality of the distribution is verified through the Shapiro-Wilk test. Comparisons for each experimental pair are indicated in the lower part of each section, showing the obtained p-value and the test used to extract the aforementioned result. Criteria for test selection are indicated in the Material and Methods section.

| Focal Adhesions Area Analysis |  |  |  |
| --- | --- | --- | --- |
|  | 0.2 kPa | 10 kPa | 27 kPa |
| Shapiro-Wilk Test (p) | 4.83e-10 | 5.8e-07 | 7.21e-10 |
| Condition | p-value | Test used | Sig |
| 0.2 kPa vs 10 kPa | 4.31e-08 | Mann-Whitney U-Test | *** |
| 0.2 kPa vs 27 kPa | 0 | Mann-Whitney U-Test | *** |
| 10 kPa vs 27 kPa | 2.47e-10 | Mann-Whitney U-Test | *** |

| Cell velocity Analysis |  |  |  |
| --- | --- | --- | --- |
|  | 0.2 kPa | 10 kPa | 27 kPa |
| Shapiro-Wilk Test (p) | 2.55e-04 | 0.00526 | 0.01420 |
| Condition | p-value | Test used | Sig |
| 0.2 kPa vs 10 kPa | 0 | Mann-Whitney U-Test | *** |
| 0.2 kPa vs 27 kPa | 7.41e-05 | Mann-Whitney U-Test | *** |
| 10 kPa vs 27 kPa | 0 | Mann-Whitney U-Test | *** |

| Western Blot for YAP/GAPDH Analysis (FC vs 0.2) |  |  |  |
| --- | --- | --- | --- |
|  | 0.2 kPa | 10 kPa | 27 kPa |
| Shapiro-Wilk Test (p) | - | 0.37380 | 0.96600 |
| Condition | p-value | Test used | Sig |
| 0.2 kPa vs 10 kPa | 0.29110 | Welch correction T-test | n.s. |
| 0.2 kPa vs 27 kPa | 0.04720 | Welch correction T-test | * |
| 10 kPa vs 27 kPa | 0.19730 | Pooled variance T-test | n.s. |

| Atomic Force Microscopy Analysis |  |  |  |
| --- | --- | --- | --- |
|  | 0.2 kPa | 10 kPa | 27 kPa |
| Shapiro-Wilk Test (p) | 0.93630 | 0.12600 | 0.60200 |
| Condition | p-value | Test used | Sig |
| 0.2 kPa vs 10 kPa | 7.73e-06 | Welch correction T-test | *** |
| 0.2 kPa vs 27 kPa | 6.66e-06 | Welch correction T-test | *** |
| 10 kPa vs 27 kPa | 0.00315 | Welch correction T-test | ** |

| YAP Nuclear translocation Analysis |  |  |  |
| --- | --- | --- | --- |
|  | 0.2 kPa | 10 kPa | 27 kPa |
| Shapiro-Wilk Test (p) | 0.95060 | 0.04831 | 0.87910 |
| Condition | p-value | Test used | Sig |
| 0.2 kPa vs 10 kPa | 1.47e-05 | Mann-Whitney U-Test | *** |
| 0.2 kPa vs 27 kPa | 1.84e-08 | Welch correction T-test | *** |
| 10 kPa vs 27 kPa | 1.55e-04 | Mann-Whitney U-Test | *** |

| EdU-positive cells Analysis |  |  |  |
| --- | --- | --- | --- |
|  | 0.2 kPa | 10 kPa | 27 kPa |
| Shapiro-Wilk Test (p) | 1.00000 | 1.00000 | 0.13050 |
| Condition | p-value | Test used | Sig |
| 0.2 kPa vs 10 kPa | 0.04290 | Pooled variance T-test | * |
| 0.2 kPa vs 27 kPa | 0.03190 | Pooled variance T-test | * |
| 10 kPa vs 27 kPa | 0.29410 | Pooled variance T-test | n.s. |

**Supplementary Table 6.** Complete statistical analysis for PANC-1-related data shown in Supplementary Figure 1. For each condition in all the experimental datasets produced (0.2 kPa, 10 kPa and 27 kPa) the normality of the distribution is verified through the Shapiro-Wilk test. Comparisons for each experimental pair are indicated in the lower part of each section, showing the obtained p-value and the test used to extract the aforementioned results. Criteria for test selection are indicated in the Material and Methods section.

| Spreading Area Analysis |  |  |  | Growth rate Analysis |  |  |  |
| --- | --- | --- | --- | --- | --- | --- | --- |
|  | 0.2 kPa | 10 kPa | 27 kPa |  | 0.2 kPa | 10 kPa | 27 kPa |
| Shapiro-Wilk Test (p) | 0.17410 | 0.09821 | 0.26180 | Shapiro-Wilk Test (p) | 0.15340 | 0.19310 | 0.55610 |
| Condition | p-value | Test used | Sig | Condition | p-value | Test used | Sig |
| 0.2 kPa vs 10 kPa | 2.43e-06 | Welch correction T-test | *** | 0.2 kPa vs 10 kPa | 8.02e-05 | Welch correction T-test | *** |
| 0.2 kPa vs 27 kPa | 6.37e-07 | Welch correction T-test | *** | 0.2 kPa vs 27 kPa | 0.00520 | Welch correction T-test | ** |
| 10 kPa vs 27 kPa | 0.01008 | Welch correction T-test | ** | 10 kPa vs 27 kPa | 1.23e-07 | Welch correction T-test | *** |

**Supplementary Table 7.** Complete statistical analysis for PANC-1-related data shown in Figure 2. For each condition in all the experimental datasets produced (0.2 kPa untreated, 27 kPa untreated, 27 kPa treated with budesonide 20 μM and 27 kPa treated with dexamethasone 20 μM) the normality of the distribution is verified through the Shapiro-Wilk test. Comparisons for each experimental pair are indicated in the lower part of each section, showing the obtained p-value and the test used to extract the aforementioned results. Criteria for test selection are indicated in the Material and Methods section.

| Focal Adhesions Area Analysis |  |  |  |  |
| --- | --- | --- | --- | --- |
|  | 0.2 kPa-NT | 27 kPa-NT | 27 kPa-B20 | 27 kPa-D20 |
| Shapiro-Wilk Test (p) | 4.87e-10 | 7.21e-12 | 2.16e-11 | 5.55e-15 |
| Condition | p-value | Test used | Sig |  |
| 0.2 kPa - NT vs 27 kPa - NT | 0 | Mann-Whitney U-Test | *** |  |
| 0.2 kPa - NT vs 27 kPa - B20 | 2.18e-06 | Mann-Whitney U-Test | *** |  |
| 0.2 kPa - NT vs 27 kPa - D20 | 1.31e-10 | Mann-Whitney U-Test | *** |  |
| 27 kPa - NT vs 27 kPa - B20 | 0 | Mann-Whitney U-Test | *** |  |
| 27 kPa - NT vs 27 kPa - D20 | 3.45e-10 | Mann-Whitney U-Test | *** |  |
| 27 kPa - B20 vs 27 kPa - D20 | 7.53e-04 | Mann-Whitney U-Test | *** |  |
| Atomic Force Microscopy Analysis |  |  |  |  |
|  | 0.2 kPa-NT | 27 kPa-NT | 27 kPa-B20 | 27 kPa-D20 |
| Shapiro-Wilk Test (p) | 0.07571 | 0.10700 | 0.02584 | 0.08282 |
| Condition | p-value | Test used | Sig |  |
| 0.2 kPa - NT vs 27 kPa - NT | 2.03e-04 | Welch correction T-test | *** |  |
| 0.2 kPa - NT vs 27 kPa - B20 | 4.12e-06 | Mann-Whitney U-Test | *** |  |
| 0.2 kPa - NT vs 27 kPa - D20 | 2.03e-05 | Welch correction T-test | *** |  |
| 27 kPa - NT vs 27 kPa - B20 | 0.03877 | Mann-Whitney U-Test | * |  |
| 27 kPa - NT vs 27 kPa - D20 | 0.06897 | Welch correction T-test | n.s. |  |
| 27 kPa - B20 vs 27 kPa - D20 | 0.37610 | Mann-Whitney U-Test | n.s. |  |
| Cell velocity Analysis |  |  |  |  |
|  | 0.2 kPa-NT | 27 kPa-NT | 27 kPa-B20 | 27 kPa-D20 |
| Shapiro-Wilk Test (p) | 0.05967 | 0.01756 | 0.06570 | 0.64370 |
| Condition | p-value | Test used | Sig |  |
| 0.2 kPa - NT vs 27 kPa - NT | 0 | Mann-Whitney U-Test | *** |  |
| 0.2 kPa - NT vs 27 kPa - B20 | 8.4e-09 | Pooled variance T-test | *** |  |
| 0.2 kPa - NT vs 27 kPa - D20 | 6.63e-23 | Pooled variance T-test | *** |  |
| 27 kPa - NT vs 27 kPa - B20 | 1.41e-14 | Mann-Whitney U-Test | *** |  |
| 27 kPa - NT vs 27 kPa - D20 | 0.01016 | Mann-Whitney U-Test | * |  |
| 27 kPa - B20 vs 27 kPa - D20 | 1.28e-09 | Welch correction T-test | *** |  |
| YAP Nuclear translocation Analysis |  |  |  |  |
|  | 0.2 kPa-NT | 27 kPa-NT | 27 kPa-B20 | 27 kPa-D20 |
| Shapiro-Wilk Test (p) | 0.86810 | 0.20440 | 0.12340 | 0.01577 |
| Condition | p-value | Test used | Sig |  |
| 0.2 kPa - NT vs 27 kPa - NT | 1.01e-05 | Welch correction T-test | *** |  |
| 0.2 kPa - NT vs 27 kPa - B20 | 0.00276 | Welch correction T-test | ** |  |
| 0.2 kPa - NT vs 27 kPa - D20 | 1.76e-06 | Mann-Whitney U-Test | *** |  |
| 27 kPa - NT vs 27 kPa - B20 | 0.00300 | Welch correction T-test | ** |  |
| 27 kPa - NT vs 27 kPa - D20 | 0.99100 | Mann-Whitney U-Test | n.s. |  |
| 27 kPa - B20 vs 27 kPa - D20 | 8.65e-05 | Mann-Whitney U-Test | *** |  |
| Western Blot for YAP/GAPDH Analysis (FC vs 27 kPa-NT) |  |  |  |  |
|  | 27 kPa-NT | 27 kPa-B20 | 27 kPa-D20 |  |
| Shapiro-Wilk Test (p) | - | 0.77880 | 0.20380 |  |
| Condition | p-value | Test used | Sig |  |
| 27 kPa - NT vs 27 kPa - B20 | 0.34810 | Welch correction T-test | n.s |  |
| 27 kPa - NT vs 27 kPa - D20 | 0.24240 | Welch correction T-test | n.s |  |
| 27 kPa - B20 vs 27 kPa - D20 | 0.87680 | Pooled Variance T-test | n.s. |  |

| EdU-positive cells Analysis |  |  |  |  |
| --- | --- | --- | --- | --- |
|  | 0.2 kPa-NT | 27 kPa-NT | 27 kPa-B20 | 27 kPa-D20 |
| Shapiro-Wilk Test (p) | 1.00000 | 0.13050 | 1.00000 | 1.00000 |
| Condition | p-value | Test used | Sig |  |
| 0.2 kPa - NT vs 27 kPa - NT | 0.03190 | Pooled Variance T-test | * |  |
| 0.2 kPa - NT vs 27 kPa - B20 | 0.27090 | Pooled Variance T-test | n.s. |  |
| 0.2 kPa - NT vs 27 kPa - D20 | 0.02350 | Pooled Variance T-test | * |  |
| 27 kPa - NT vs 27 kPa - B20 | 0.02461 | Pooled Variance T-test | * |  |
| 27 kPa - NT vs 27 kPa - D20 | 0.58520 | Pooled Variance T-test | n.s. |  |
| 27 kPa - B20 vs 27 kPa - D20 | 0.03388 | Pooled Variance T-test | * |  |

**Supplementary Table 8.** Complete statistical analysis for PANC-1-related data shown in Supplementary Figure 2. For each condition in all the experimental datasets produced (0.2 kPa untreated, 27 kPa untreated, 27 kPa treated with budesonide 20 μM and 27 kPa treated with dexamethasone 20 μM) the normality of the distribution is verified through the Shapiro-Wilk test. Comparisons for each experimental pair are indicated in the lower part of each section, showing the obtained p-value and the test used to extract the aforementioned results. Criteria for test selection are indicated in Material and Methods section.

| Spreading Area Analysis |  |  |  |  |
| --- | --- | --- | --- | --- |
|  | 0.2 kPa-NT | 27 kPa-NT | 27 kPa-B20 | 27 kPa-D20 |
| Shapiro-Wilk Test (p) | 0.27530 | 0.67900 | 0.00164 | 0.00551 |
| Condition | p-value | Test used | Sig |  |
| 0.2 kPa - NT vs 27 kPa - NT | 5.7e-14 | Welch correction T-test | *** |  |
| 0.2 kPa - NT vs 27 kPa - B20 | 2.08e-05 | Mann-Whitney U-Test | *** |  |
| 0.2 kPa - NT vs 27 kPa - D20 | 7.88e-11 | Mann-Whitney U-Test | *** |  |
| 27 kPa - NT vs 27 kPa - B20 | 0.00173 | Mann-Whitney U-Test | ** |  |
| 27 kPa - NT vs 27 kPa - D20 | 0.33110 | Mann-Whitney U-Test | n.s. |  |
| 27 kPa - B20 vs 27 kPa - D20 | 0.00564 | Mann-Whitney U-Test | ** |  |
| Growth rate Analysis |  |  |  |  |
|  | 0.2 kPa-NT | 27 kPa-NT | 27 kPa-B20 | 27 kPa-D20 |
| Shapiro-Wilk Test (p) | 0.02960 | 0.23960 | 0.36980 | 0.41260 |
| Condition | p-value | Test used | Sig |  |
| 0.2 kPa - NT vs 27 kPa - NT | 3.28e-06 | Mann-Whitney U-Test | *** |  |
| 0.2 kPa - NT vs 27 kPa - B20 | 0.40170 | Mann-Whitney U-Test | n.s. |  |
| 0.2 kPa - NT vs 27 kPa - D20 | 1.14e-04 | Mann-Whitney U-Test | *** |  |
| 27 kPa - NT vs 27 kPa - B20 | 2.63e-07 | Pooled variance T-test | *** |  |
| 27 kPa - NT vs 27 kPa - D20 | 0.01802 | Pooled variance T-test | * |  |
| 27 kPa - B20 vs 27 kPa - D20 | 6.06e-05 | Mann-Whitney U-Test | *** |  |

**Supplementary Table 9.**
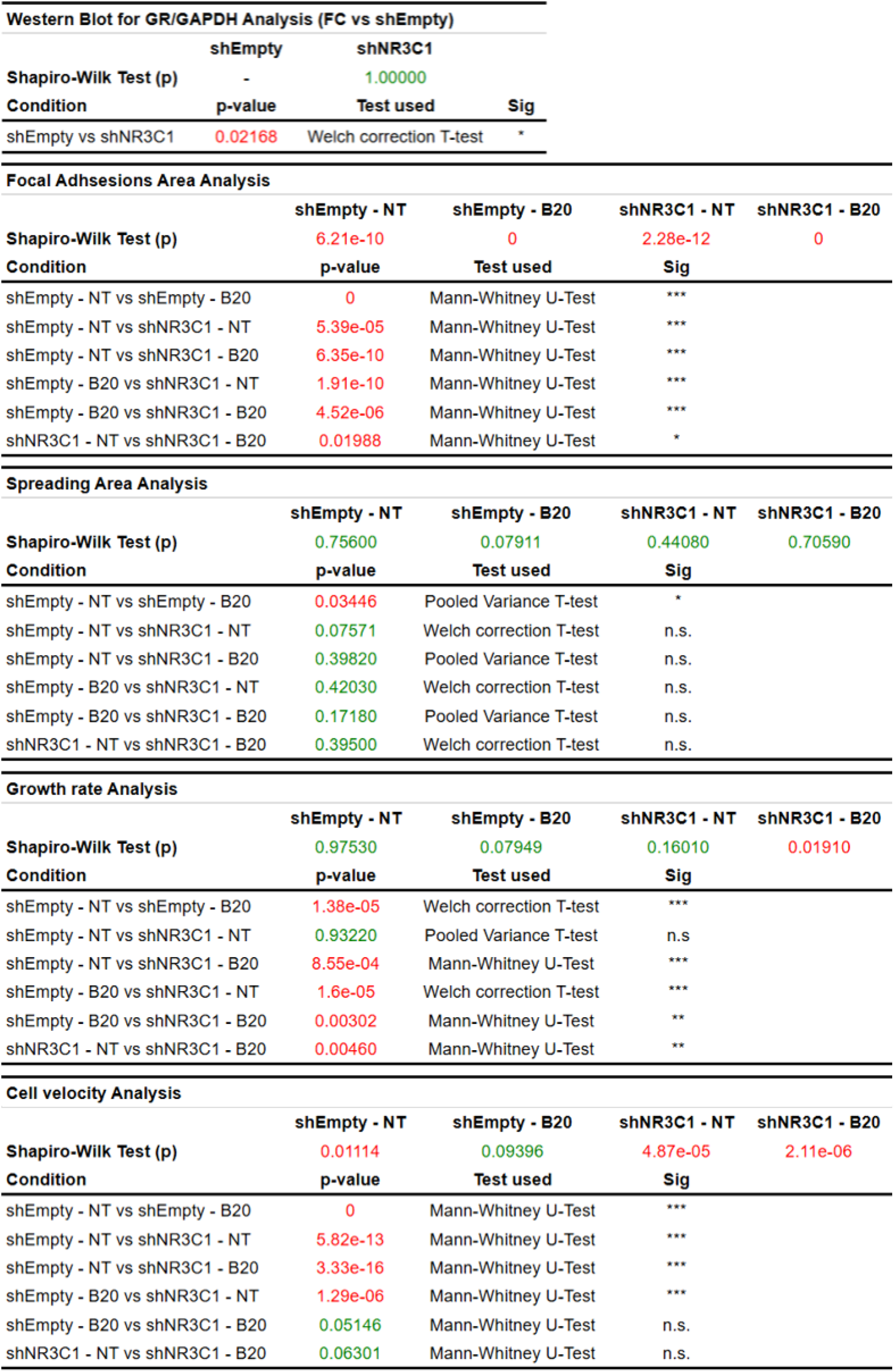
Complete statistical analysis for NR3C1-silenced PANC-1-related data shown in Supplementary Figure 3. For each condition in all the experimental datasets produced (*^shEmpty^*PANC-1 untreated, *^shEmpty^*PANC-1 treated with budesonide 20 µM, *^shNR3C1^*PANC-1 untreated and *^shNR3C1^*PANC-1 treated with budesonide 20 µM) the normality of the distribution is verified through the Shapiro-Wilk test. Comparisons for each experimental pair are indicated in the lower part of each section, showing the obtained pvalue and the test used to extract the aforementioned results. Criteria for test selection are indicated in Material and Methods section.

**Supplementary Table 10.** Complete statistical analysis for PDAC#253-related data shown in Figure 3. For each condition in all the experimental datasets produced (0.2 kPa untreated, 27 kPa untreated, 27 kPa treated with budesonide 20 µM and 27 kPa treated with dexamethasone 20 µM) the normality of the distribution is verified through the Shapiro-Wilk test. Comparisons for each experimental pair are indicated in the lower part of each section, showing the obtained p-value and the test used to extract the aforementioned results. Criteria for test selection are indicated in the Material and Methods section.

| Focal Adhesions Area Analysis |  |  |  |  |
| --- | --- | --- | --- | --- |
|  | 0.2 kPa-NT | 27 kPa-NT | 27 kPa-B20 | 27 kPa-D20 |
| Shapiro-Wilk Test (p) | 3.16e-12 | 0 | 0 | 0 |
| Condition | p-value | Test used | Sig |  |
| 0.2 kPa - NT vs 27 kPa - NT | 0 | Mann-Whitney U-Test | *** |  |
| 0.2 kPa - NT vs 27 kPa - B20 | 0.02319 | Mann-Whitney U-Test | * |  |
| 0.2 kPa - NT vs 27 kPa - D20 | 0 | Mann-Whitney U-Test | *** |  |
| 27 kPa - NT vs 27 kPa - B20 | 0 | Mann-Whitney U-Test | *** |  |
| 27 kPa - NT vs 27 kPa - D20 | 0.07238 | Mann-Whitney U-Test | n.s. |  |
| 27 kPa - B20 vs 27 kPa - D20 | 0 | Mann-Whitney U-Test | *** |  |
| Cell velocity Analysis |  |  |  |  |
|  | 0.2 kPa-NT | 27 kPa-NT | 27 kPa-B20 | 27 kPa-D20 |
| Shapiro-Wilk Test (p) | 1.76e-05 | 0.52070 | 2.48e-05 | 0.28310 |
| Condition | p-value | Test used | Sig |  |
| 0.2 kPa - NT vs 27 kPa - NT | 0 | Mann-Whitney U-Test | *** |  |
| 0.2 kPa - NT vs 27 kPa - B20 | 0.09582 | Mann-Whitney U-Test | n.s. |  |
| 0.2 kPa - NT vs 27 kPa - D20 | 0 | Mann-Whitney U-Test | *** |  |
| 27 kPa - NT vs 27 kPa - B20 | 0 | Mann-Whitney U-Test | *** |  |
| 27 kPa - NT vs 27 kPa - D20 | 1.58e-10 | Pooled Variance T-test | *** |  |
| 27 kPa - B20 vs 27 kPa - D20 | 0 | Mann-Whitney U-Test | *** |  |
| YAP Nuclear translocation Analysis |  |  |  |  |
|  | 0.2 kPa-NT | 27 kPa-NT | 27 kPa-B20 | 27 kPa-D20 |
| Shapiro-Wilk Test (p) | 0.72430 | 0.71550 | 0.03613 | 0.50740 |
| Condition | p-value | Test used | Sig |  |
| 0.2 kPa - NT vs 27 kPa - NT | 5.98e-23 | Pooled Variance T-test | *** |  |
| 0.2 kPa - NT vs 27 kPa - B20 | 0.46370 | Mann-Whitney U-Test | n.s. |  |
| 0.2 kPa - NT vs 27 kPa - D20 | 4.99e-13 | Welch correction T-test | *** |  |
| 27 kPa - NT vs 27 kPa - B20 | 5.45e-12 | Mann-Whitney U-Test | *** |  |
| 27 kPa - NT vs 27 kPa - D20 | 0.80510 | Welch correction T-test | n.s. |  |
| 27 kPa - B20 vs 27 kPa - D20 | 1.25e-08 | Mann-Whitney U-Test | *** |  |

**Supplementary Table 11.** Complete statistical analysis for PDAC#253-related data shown in Supplementary Figure 4. For each condition in all the experimental datasets produced (0.2 kPa untreated, 27 kPa untreated, 27 kPa treated with budesonide 20 µM and 27 kPa treated with dexamethasone 20 µM) the normality of the distribution is verified through the Shapiro-Wilk test. Comparisons for each experimental pair are indicated in the lower part of each section, showing the obtained p-value and the test used to extract the aforementioned results. Criteria for test selection are indicated in the Material and Methods section.

| Spreading Area Analysis |  |  |  |  |
| --- | --- | --- | --- | --- |
|  | 0.2 kPa-NT | 27 kPa-NT | 27 kPa-B20 | 27 kPa-D20 |
| Shapiro-Wilk Test (p) | 0.00258 | 7.11e-05 | 0.00451 | 0.01369 |
| Condition | p-value | Test used | Sig |  |
| 0.2 kPa - NT vs 27 kPa - NT | 2.67e-15 | Mann-Whitney U-Test | *** |  |
| 0.2 kPa - NT vs 27 kPa - B20 | 0.06618 | Mann-Whitney U-Test | n.s. |  |
| 0.2 kPa - NT vs 27 kPa - D20 | 0 | Mann-Whitney U-Test | *** |  |
| 27 kPa - NT vs 27 kPa - B20 | 7.25e-10 | Mann-Whitney U-Test | *** |  |
| 27 kPa - NT vs 27 kPa - D20 | 8.22e-15 | Mann-Whitney U-Test | *** |  |
| 27 kPa - B20 vs 27 kPa - D20 | 0 | Mann-Whitney U-Test | *** |  |
| Growth rate Analysis |  |  |  |  |
|  | 0.2 kPa-NT | 27 kPa-NT | 27 kPa-B20 | 27 kPa-D20 |
| Shapiro-Wilk Test (p) | 0.28290 | 0.02909 | 0.10020 | 0.18430 |
| Condition | p-value | Test used | Sig |  |
| 0.2 kPa - NT vs 27 kPa - NT | 0.38350 | Mann-Whitney U-Test | n.s. |  |
| 0.2 kPa - NT vs 27 kPa - B20 | 6.98e-04 | Welch correction T-test | *** |  |
| 0.2 kPa - NT vs 27 kPa - D20 | 0.22180 | Pooled Variance T-test | n.s. |  |
| 27 kPa - NT vs 27 kPa - B20 | 1.39e-04 | Mann-Whitney U-Test | *** |  |
| 27 kPa - NT vs 27 kPa - D20 | 0.06975 | Mann-Whitney U-Test | n.s. |  |
| 27 kPa - B20 vs 27 kPa - D20 | 2.14e-04 | Pooled Variance T-test | *** |  |

**Supplementary Table 12.** Complete statistical analysis for PDAC#354-related data shown in Supplementary Figure 4. For each condition in all the experimental datasets produced (0.2 kPa untreated, 27 kPa untreated, 27 kPa treated with budesonide 20 µM and 27 kPa treated with dexamethasone 20 µM) the normality of the distribution is verified through the Shapiro-Wilk test. Comparisons for each experimental pair are indicated in the lower part of each section, showing the obtained p-value and the test used to extract the aforementioned results. Criteria for test selection are indicated in the Material and Methods section.

| Spreading Area Analysis |  |  |  |  |
| --- | --- | --- | --- | --- |
|  | 0.2 kPa-NT | 27 kPa-NT | 27 kPa-B20 | 27 kPa-D20 |
| Shapiro-Wilk Test (p) | 0.00644 | 0.00822 | 0.72610 | 0.00144 |
| Condition | p-value | Test used | Sig |  |
| 0.2 kPa - NT vs 27 kPa - NT | 4.25e-12 | Mann-Whitney U-Test | *** |  |
| 0.2 kPa - NT vs 27 kPa - B20 | 2e-06 | Mann-Whitney U-Test | *** |  |
| 0.2 kPa - NT vs 27 kPa - D20 | 6.29e-13 | Mann-Whitney U-Test | *** |  |
| 27 kPa - NT vs 27 kPa - B20 | 3.75e-05 | Mann-Whitney U-Test | *** |  |
| 27 kPa - NT vs 27 kPa - D20 | 0.92240 | Mann-Whitney U-Test | n.s. |  |
| 27 kPa - B20 vs 27 kPa - D20 | 4.74e-05 | Mann-Whitney U-Test | *** |  |
| Growth rate Analysis |  |  |  |  |
|  | 0.2 kPa-NT | 27 kPa-NT | 27 kPa-B20 | 27 kPa-D20 |
| Shapiro-Wilk Test (p) | 0.79180 | 0.02709 | 0.87150 | 0.96570 |
| Condition | p-value | Test used | Sig |  |
| 0.2 kPa - NT vs 27 kPa - NT | 0.00791 | Mann-Whitney U-Test | ** |  |
| 0.2 kPa - NT vs 27 kPa - B20 | 1.21e-13 | Pooled Variance T-test | *** |  |
| 0.2 kPa - NT vs 27 kPa - D20 | 6.76e-09 | Pooled Variance T-test | *** |  |
| 27 kPa - NT vs 27 kPa - B20 | 6.28e-08 | Mann-Whitney U-Test | *** |  |
| 27 kPa - NT vs 27 kPa - D20 | 1.42e-07 | Mann-Whitney U-Test | *** |  |
| 27 kPa - B20 vs 27 kPa - D20 | 1.56e-06 | Welch correction T-test | *** |  |
| Cell velocity Analysis |  |  |  |  |
|  | 0.2 kPa-NT | 27 kPa-NT | 27 kPa-B20 | 27 kPa-D20 |
| Shapiro-Wilk Test (p) | 0.02321 | 0.03946 | 6.26e-05 | 0.47000 |
| Condition | p-value | Test used | Sig |  |
| 0.2 kPa - NT vs 27 kPa - NT | 0 | Mann-Whitney U-Test | *** |  |
| 0.2 kPa - NT vs 27 kPa - B20 | 6.88e-07 | Mann-Whitney U-Test | *** |  |
| 0.2 kPa - NT vs 27 kPa - D20 | 0 | Mann-Whitney U-Test | *** |  |
| 27 kPa - NT vs 27 kPa - B20 | 0 | Mann-Whitney U-Test | *** |  |
| 27 kPa - NT vs 27 kPa - D20 | 9.25e-14 | Mann-Whitney U-Test | *** |  |
| 27 kPa - B20 vs 27 kPa - D20 | 8.71e-10 | Mann-Whitney U-Test | *** |  |
| YAP Nuclear translocation Analysis |  |  |  |  |
|  | 0.2 kPa-NT | 27 kPa-NT | 27 kPa-B20 | 27 kPa-D20 |
| Shapiro-Wilk Test (p) | 0.02293 | 0.55380 | 0.63610 | 0.41110 |
| Condition | p-value | Test used | Sig |  |
| 0.2 kPa - NT vs 27 kPa - NT | 1.31e-13 | Mann-Whitney U-Test | *** |  |
| 0.2 kPa - NT vs 27 kPa - B20 | 0.18180 | Mann-Whitney U-Test | n.s. |  |
| 0.2 kPa - NT vs 27 kPa - D20 | 7.48e-08 | Mann-Whitney U-Test | *** |  |
| 27 kPa - NT vs 27 kPa - B20 | 1.58e-12 | Welch correction T-test | *** |  |
| 27 kPa - NT vs 27 kPa - D20 | 2.02e-05 | Welch correction T-test | *** |  |
| 27 kPa - B20 vs 27 kPa - D20 | 1.46e-05 | Pooled Variance T-test | *** |  |

**Supplementary Table 13.** Complete statistical analysis for PANC-1 nuclear and nucleoskeletal morphometry data shown in Figure 5 and Supplementary Figure 6. For each condition in all the experimental datasets produced (0.2 kPa untreated, 27 kPa untreated and 27 kPa treated with budesonide 20μM) the normality of the distribution is verified through the Shapiro-Wilk test. Comparisons for each experimental pair are indicated in the lower part of each section, showing the obtained p-value and the test used to extract the aforementioned results. Criteria for test selection are indicated in the Material and Methods section.

|  |  |  |  |
| --- | --- | --- | --- |
| **Lamin A/C Intensity Analysis** |  |  |  |
|  | **0.2 kPa-NT** | **27 kPa-NT** | **27 kPa-B20** |
| **Shapiro-Wilk Test (p)** | **0.30460** | **0.00973** | **0.75180** |
| **Condition** | **p-value** | **Test used** | **Sig** |
| 0.2 kPa - NT vs 27 kPa - NT | 0.00258 | Mann-Whitney U-Test | ** |
| 0.2 kPa - NT vs 27 kPa - B20 | 4.26e-04 | Welch correction T-test | *** |
| 27 kPa - NT vs 27 kPa - B20 | 1.4e-08 | Mann-Whitney U-Test | *** |
| **Lamin B1 Intensity Analysis** |  |  |  |
|  | **0.2 kPa-NT** | **27 kPa-NT** | **27 kPa-B20** |
| **Shapiro-Wilk Test (p)** | **0.01644** | **0.08926** | **0.69870** |
| **Condition** | **p-value** | **Test used** | **Sig** |
| 0.2 kPa - NT vs 27 kPa - NT | 6.83e-13 | Mann-Whitney U-Test | *** |
| 0.2 kPa - NT vs 27 kPa - B20 | 4.19e-09 | Mann-Whitney U-Test | *** |
| 27 kPa - NT vs 27 kPa - B20 | 0.00157 | Pooled variance T-test | ** |
| **Wrinkling Index Analysis** |  |  |  |
|  | **0.2 kPa-NT** | **27 kPa-NT** | **27 kPa-B20** |
| **Shapiro-Wilk Test (p)** | **0.07064** | **0.21020** | **0.01845** |
| **Condition** | **p-value** | **Test used** | **Sig** |
| 0.2 kPa - NT vs 27 kPa - NT | 0 | Pooled variance T-test | *** |
| 0.2 kPa - NT vs 27 kPa - B20 | 1.14e-04 | Mann-Whitney U-Test | *** |
| 27 kPa - NT vs 27 kPa - B20 | 5.78e-13 | Mann-Whitney U-Test | *** |
| **Nuclear Volume Analysis** |  |  |  |
|  | **0.2 kPa-NT** | **27 kPa-NT** | **27 kPa-B20** |
| **Shapiro-Wilk Test (p)** | **0.01756** | **0.00465** | **0.05050** |
| **Condition** | **p-value** | **Test used** | **Sig** |
| 0.2 kPa - NT vs 27 kPa - NT | 0.01355 | Mann-Whitney U-Test | ** |
| 0.2 kPa - NT vs 27 kPa - B20 | 0.09415 | Mann-Whitney U-Test | n.s. |
| 27 kPa - NT vs 27 kPa - B20 | 1.64e-04 | Mann-Whitney U-Test | *** |
| **Nuclear Height Analysis** |  |  |  |
|  | **0.2 kPa-NT** | **27 kPa-NT** | **27 kPa-B20** |
| **Shapiro-Wilk Test (p)** | **0.07764** | **0.98770** | **0.01163** |
| **Condition** | **p-value** | **Test used** | **Sig** |
| 0.2 kPa - NT vs 27 kPa - NT | 0.85270 | Pooled Variance T-test | n.s. |
| 0.2 kPa - NT vs 27 kPa - B20 | 2.05e-06 | Mann-Whitney U-Test | *** |
| 27 kPa - NT vs 27 kPa - B20 | 1.14e-16 | Mann-Whitney U-Test | *** |
| **Major Axis of Nuclear elliptical section Analysis** |  |  |  |
|  | **0.2 kPa-NT** | **27 kPa-NT** | **27 kPa-B20** |
| **Shapiro-Wilk Test (p)** | **0.16140** | **0.04367** | **0.15590** |
| **Condition** | **p-value** | **Test used** | **Sig** |
| 0.2 kPa - NT vs 27 kPa - NT | 0.01320 | Mann-Whitney U-Test | * |
| 0.2 kPa - NT vs 27 kPa - B20 | 0.86440 | Pooled Variance T-test | n.s. |
| 27 kPa - NT vs 27 kPa - B20 | 0.01171 | Mann-Whitney U-Test | * |
| **Minor Axis of Nuclear elliptical section Analysis** |  |  |  |
|  | **0.2 kPa-NT** | **27 kPa-NT** | **27 kPa-B20** |
| **Shapiro-Wilk Test (p)** | **0.82060** | **0.01974** | **0.11640** |
| **Condition** | **p-value** | **Test used** | **Sig** |
| 0.2 kPa - NT vs 27 kPa - NT | 0.10350 | Mann-Whitney U-Test | n.s. |
| 0.2 kPa - NT vs 27 kPa - B20 | 0.89520 | Pooled Variance T-test | n.s. |
| 27 kPa - NT vs 27 kPa - B20 | 0.09154 | Mann-Whitney U-Test | n.s. |

**Supplementary Table 14.** Complete statistical analysis for PDAC#253 nuclear and nucleoskeletal morphometry data shown in Figure 5 and Supplementary Figure 6. For each condition in all the experimental datasets produced (0.2 kPa untreated, 27 kPa untreated and 27 kPa treated with budesonide 20μM) the normality of the distribution is verified through the Shapiro-Wilk test. Comparisons for each experimental pair are indicated in the lower part of each section, showing the obtained p-value and the test used to extract the aforementioned result. Criteria for test selection are indicated in the Material and Methods section.

| Lamin A/C Intensity Analysis |  |  |  |
| --- | --- | --- | --- |
|  | 0.2 kPa-NT | 27 kPa-NT | 27 kPa-B20 |
| Shapiro-Wilk Test (p) | 0.83080 | 0.03124 | 0.28360 |
| Condition | p-value | Test used | Sig |
| 0.2 kPa - NT vs 27 kPa - NT | 9.13e-11 | Mann-Whitney U-Test | *** |
| 0.2 kPa - NT vs 27 kPa - B20 | 2.22e-13 | Pooled variance T-test | *** |
| 27 kPa - NT vs 27 kPa - B20 | 2.2e-10 | Mann-Whitney U-Test | *** |

| Lamin B1 Intensity Analysis |  |  |  |
| --- | --- | --- | --- |
|  | 0.2 kPa-NT | 27 kPa-NT | 27 kPa-B20 |
| Shapiro-Wilk Test (p) | 0.02842 | 0.57760 | 0.15290 |
| Condition | p-value | Test used | Sig |
| 0.2 kPa - NT vs 27 kPa - NT | 1.98e-10 | Mann-Whitney U-Test | *** |
| 0.2 kPa - NT vs 27 kPa - B20 | 1.38e-12 | Mann-Whitney U-Test | *** |
| 27 kPa - NT vs 27 kPa - B20 | 2.15e-04 | Pooled variance T-test | *** |

| Wrinkling Index Analysis |  |  |  |
| --- | --- | --- | --- |
|  | 0.2 kPa-NT | 27 kPa-NT | 27 kPa-B20 |
| Shapiro-Wilk Test (p) | 0.54490 | 0.22570 | 0.01034 |
| Condition | p-value | Test used | Sig |
| 0.2 kPa - NT vs 27 kPa - NT | 2.3e-04 | Pooled variance T-test | *** |
| 0.2 kPa - NT vs 27 kPa - B20 | 0.03863 | Mann-Whitney U-Test | * |
| 27 kPa - NT vs 27 kPa - B20 | 5.9e-07 | Mann-Whitney U-Test | *** |

| Nuclear Volume Analysis |  |  |  |
| --- | --- | --- | --- |
|  | 0.2 kPa-NT | 27 kPa-NT | 27 kPa-B20 |
| Shapiro-Wilk Test (p) | 0.26710 | 0.04756 | 0.16390 |
| Condition | p-value | Test used | Sig |
| 0.2 kPa - NT vs 27 kPa - NT | 2.07e-11 | Mann-Whitney U-Test | *** |
| 0.2 kPa - NT vs 27 kPa - B20 | 0.01326 | Pooled Variance T-test | * |
| 27 kPa - NT vs 27 kPa - B20 | 7.77e-16 | Mann-Whitney U-Test | *** |

| Nuclear Height Analysis |  |  |  |
| --- | --- | --- | --- |
|  | 0.2 kPa-NT | 27 kPa-NT | 27 kPa-B20 |
| Shapiro-Wilk Test (p) | 0.77250 | 0.33700 | 0.14920 |
| Condition | p-value | Test used | Sig |
| 0.2 kPa - NT vs 27 kPa - NT | 6.18e-04 | Welch correction T-test | *** |
| 0.2 kPa - NT vs 27 kPa - B20 | 2.22e-16 | Pooled Variance T-test | *** |
| 27 kPa - NT vs 27 kPa - B20 | 0 | Welch correction T-test | *** |

| Major Axis of Nuclear elliptical section Analysis |  |  |  |
| --- | --- | --- | --- |
|  | 0.2 kPa-NT | 27 kPa-NT | 27 kPa-B20 |
| Shapiro-Wilk Test (p) | 0.57170 | 0.47050 | 0.02942 |
| Condition | p-value | Test used | Sig |
| 0.2 kPa - NT vs 27 kPa - NT | 2.67e-08 | Welch correction T-test | *** |
| 0.2 kPa - NT vs 27 kPa - B20 | 9.15e-05 | Mann-Whitney U-Test | *** |
| 27 kPa - NT vs 27 kPa - B20 | 0.13830 | Mann-Whitney U-Test | n.s. |

| Minor Axis of Nuclear elliptical section Analysis |  |  |  |
| --- | --- | --- | --- |
|  | 0.2 kPa-NT | 27 kPa-NT | 27 kPa-B20 |
| Shapiro-Wilk Test (p) | 0.87970 | 0.01725 | 0.10400 |
| Condition | p-value | Test used | Sig |
| 0.2 kPa - NT vs 27 kPa - NT | 0.01547 | Mann-Whitney U-Test | * |
| 0.2 kPa - NT vs 27 kPa - B20 | 0.70450 | Pooled Variance T-test | n.s. |
| 27 kPa - NT vs 27 kPa - B20 | 0.01152 | Mann-Whitney U-Test | * |

**Supplementary Table 15.** Complete statistical analysis for silenced PANC-1 nuclear and nucleoskeletal morphometry data shown in Supplementary Figure 6. For each condition in all the experimental datasets produced (shEmpty-PANC-1 untreated, shEmpty-PANC-1 treated with budesonide 20 µM, shNR3C1-PANC-1 untreated and shNR3C1-PANC-1 treated with budesonide 20 µM) the normality of the distribution is verified through the Shapiro-Wilk test. Comparisons for each experimental pair are indicated in the lower part of each section, showing the obtained p-value and the test used to extract the aforementioned result. Criteria for test selection are indicated in the Material and Methods section.

| Lamin A/C Intensity Analysis |  |  |  |  |
| --- | --- | --- | --- | --- |
|  | shEmpty - NT | shEmpty - B20 | shNR3C1 - NT | shNR3C1 - B20 |
| Shapiro-Wilk Test (p) | 0.02123 | 0.14300 | 0.34710 | 0.32470 |
| Condition | p-value | Test used | Sig |  |
| shEmpty - NT vs shEmpty - B20 | 1.93e-05 | Mann-Whitney U-Test | *** |  |
| shEmpty - NT vs shNR3C1 - NT | 4.26e-07 | Mann-Whitney U-Test | *** |  |
| shEmpty - NT vs shNR3C1 - B20 | 1.3e-06 | Mann-Whitney U-Test | *** |  |
| shEmpty - B20 vs shNR3C1 - NT | 0.86940 | Pooled Variance T-test | n.s. |  |
| shEmpty - B20 vs shNR3C1 - B20 | 0.89530 | Pooled Variance T-test | n.s. |  |
| shNR3C1 - NT vs shNR3C1 - B20 | 0.73100 | Pooled Variance T-test | n.s. |  |

| Lamin B1 Intensity Analysis |  |  |  |  |
| --- | --- | --- | --- | --- |
|  | shEmpty - NT | shEmpty - B20 | shNR3C1 - NT | shNR3C1 - B20 |
| Shapiro-Wilk Test (p) | 0.03359 | 0.00428 | 0.08251 | 0.01147 |
| Condition | p-value | Test used | Sig |  |
| shEmpty - NT vs shEmpty - B20 | 0.02758 | Mann-Whitney U-Test | *** |  |
| shEmpty - NT vs shNR3C1 - NT | 0.00251 | Mann-Whitney U-Test | *** |  |
| shEmpty - NT vs shNR3C1 - B20 | 0.00141 | Mann-Whitney U-Test | *** |  |
| shEmpty - B20 vs shNR3C1 - NT | 0.12090 | Mann-Whitney U-Test | n.s. |  |
| shEmpty - B20 vs shNR3C1 - B20 | 0.10880 | Mann-Whitney U-Test | n.s. |  |
| shNR3C1 - NT vs shNR3C1 - B20 | 0.94080 | Mann-Whitney U-Test | n.s. |  |

| Wrinkling Index Analysis |  |  |  |  |
| --- | --- | --- | --- | --- |
|  | shEmpty - NT | shEmpty - B20 | shNR3C1 - NT | shNR3C1 - B20 |
| Shapiro-Wilk Test (p) | 0.07396 | 0.35850 | 0.00937 | 0.53350 |
| Condition | p-value | Test used | Sig |  |
| shEmpty - NT vs shEmpty - B20 | 2.84e-06 | Welch correction T-test | *** |  |
| shEmpty - NT vs shNR3C1 - NT | 5.05e-06 | Mann-Whitney U-Test | *** |  |
| shEmpty - NT vs shNR3C1 - B20 | 3.83e-05 | Welch correction T-test | *** |  |
| shEmpty - B20 vs shNR3C1 - NT | 0.27530 | Mann-Whitney U-Test | n.s. |  |
| shEmpty - B20 vs shNR3C1 - B20 | 0.56870 | Pooled Variance T-test | n.s. |  |
| shNR3C1 - NT vs shNR3C1 - B20 | 0.79880 | Mann-Whitney U-Test | n.s. |  |

| Nuclear Volume Analysis |  |  |  |  |
| --- | --- | --- | --- | --- |
|  | shEmpty - NT | shEmpty - B20 | shNR3C1 - NT | shNR3C1 - B20 |
| Shapiro-Wilk Test (p) | 0.00480 | 0.01239 | 0.01860 | 0.00461 |
| Condition | p-value | Test used | Sig |  |
| shEmpty - NT vs shEmpty - B20 | 1.21e-04 | Mann-Whitney U-Test | *** |  |
| shEmpty - NT vs shNR3C1 - NT | 0.86620 | Mann-Whitney U-Test | n.s. |  |
| shEmpty - NT vs shNR3C1 - B20 | 6.14e-05 | Mann-Whitney U-Test | *** |  |
| shEmpty - B20 vs shNR3C1 - NT | 5.86e-04 | Mann-Whitney U-Test | *** |  |
| shEmpty - B20 vs shNR3C1 - B20 | 0.50530 | Mann-Whitney U-Test | n.s. |  |
| shNR3C1 - NT vs shNR3C1 - B20 | 9.72e-06 | Mann-Whitney U-Test | *** |  |

| Nuclear Height Analysis |  |  |  |  |
| --- | --- | --- | --- | --- |
|  | shEmpty - NT | shEmpty - B20 | shNR3C1 - NT | shNR3C1 - B20 |
| Shapiro-Wilk Test (p) | 0.44800 | 0.24970 | 0.46050 | 0.76720 |
| Condition | p-value | Test used | Sig |  |
| shEmpty - NT vs shEmpty - B20 | 4.09e-07 | Pooled Variance T-test | *** |  |
| shEmpty - NT vs shNR3C1 - NT | 0.59370 | Pooled Variance T-test | n.s. |  |
| shEmpty - NT vs shNR3C1 - B20 | 5.38e-06 | Pooled Variance T-test | *** |  |
| shEmpty - B20 vs shNR3C1 - NT | 1.95e-04 | Welch correction T-test | *** |  |
| shEmpty - B20 vs shNR3C1 - B20 | 0.72860 | Pooled Variance T-test | n.s. |  |
| shNR3C1 - NT vs shNR3C1 - B20 | 3.35e-04 | Pooled Variance T-test | *** |  |

| Major Axis of Nuclear elliptical section Analysis |  |  |  |
| --- | --- | --- | --- |
|  | 0.2 kPa-NT | 27 kPa-NT | 27 kPa-B20 |
| Shapiro-Wilk Test (p) | 0.57170 | 0.47050 | 0.02942 |
| Condition | p-value | Test used | Sig |
| 0.2 kPa - NT vs 27 kPa - NT | 2.67e-08 | Welch correction T-test | *** |
| 0.2 kPa - NT vs 27 kPa - B20 | 9.15e-05 | Mann-Whitney U-Test | *** |
| 27 kPa - NT vs 27 kPa - B20 | 0.13830 | Mann-Whitney U-Test | n.s. |
| Minor Axis of Nuclear elliptical section Analysis |  |  |  |
|  | 0.2 kPa-NT | 27 kPa-NT | 27 kPa-B20 |
| Shapiro-Wilk Test (p) | 0.87970 | 0.01725 | 0.10400 |
| Condition | p-value | Test used | Sig |
| 0.2 kPa - NT vs 27 kPa - NT | 0.01547 | Mann-Whitney U-Test | * |
| 0.2 kPa - NT vs 27 kPa - B20 | 0.70450 | Pooled Variance T-test | n.s. |
| 27 kPa - NT vs 27 kPa - B20 | 0.01152 | Mann-Whitney U-Test | * |

